# RhoGAP54D promotes cell size asymmetry and inhibits pulsatile Myosin activity in *Drosophila* neural stem cells

**DOI:** 10.1101/2025.09.15.676233

**Authors:** Nicolas Loyer, Guozhen Li, Jens Januschke

## Abstract

The Actomyosin cortex is a highly dynamic part of the cytoskeleton and cells can modulate its properties. This is important for essential cellular functions including migration, cell shape control and cleavage furrow positioning during cell division. Asymmetrically dividing *Drosophila* neural stem cells, called neuroblasts, exhibit stereotypic patterns of Myosin dynamics along the cell cycle. Some of the Myosin dynamics manifest as flows and are important to asymmetrically position the cleavage furrow which establishes the cell size-difference between the neuroblast and its daughter cell. Myosin can be regulated by small GTPases and their GAPs and GEFs, providing spatio-temporal control. Rho kinase downstream of the small GTPase Rho regulates Myosin activity, yet the regulation and contribution of its GEFs and GAPs to asymmetric neuroblast division are not resolved. Here, we systematically analysed the localization of RhoGAPs and RhoGEFs expressed in neuroblasts and identified the ARHGAP19 homolog RhoGAP54D as a regulator of Myosin dynamics. A cytoplasmic pool of RhoGAP54D suppresses pulsatile Myosin activity during interphase and metaphase, regulated by the nucleoporin Members Only via slow nuclear import. In anaphase RhoGAP54D is recruited to the apical cortex and constitutes the trigger that initiates the directional Myosin flow that positions the cleavage furrow. This depends on PsGEF which is itself recruited downstream of Par3/Baz and Pins, linking cell polarity to RhoGAP54D activity to promote daughter cell size asymmetry during neuroblast division.

## Introduction

The Actomyosin cortex displays an array of behaviours, ranging from pulsatile contractions to polarized flows, and contractile structures driving many cellular processes such as cell shape modulation, migration and cytokinesis (Staddon et al. 2022). While some aspects of Actomyosin behaviour reflect its emergent properties, upstream regulators often provide spatio-temporal control. The small Rho GTPases of the Rho family serve as molecular switches in this context regulating processes including tissue morphogenesis and cell polarity (Riento and Ridley 2003; Jaffe and Hall 2005; Park and Bi 2007). Control of GTPase activity is achieved through regulatory proteins called guanine nucleotide exchange factors (GEFs, ‘activators’) and GTPase activating proteins (GAPs, ‘inactivators’). This regulation occurs frequently at distinct subcellular sites triggering local responses at specific time points (Cherfils and Zeghouf 2013).

Coordinated cytoskeletal remodelling occurs along the cell cycle of *Drosophila* neuroblasts that generate the nervous system of the adult fly during development. Neuroblasts divisions are asymmetric in fate and size: they generate a large self-renewed neuroblast with stem cell potential and a much smaller differentiating ganglion mother cell (GMC), (Delgado and Cabernard 2020; Loyer and Januschke 2020). This asymmetry relies on cell polarity: The PAR complex including Par3 (Bazooka, Baz in flies(Wodarz et al. 1999)) localises to the apical cortex in mitosis and drives the basal localisation of the cell fate determinants inherited by the GMC upon division (Gallaud et al. 2017). The apical Pins complex controls spindle orientation (Parmentier et al. 2000; Schaefer et al. 2000; Yu et al. 2000) via Mud/Numa (Bowman et al. 2006) as well as cell size asymmetry (Parmentier et al. 2000), the mechanism of which is not resolved. Both the localisation of polarised determinants and size asymmetry also somehow depend on Actomyosin (Broadus and Doe 1997; Lu et al. 1999; Cabernard et al. 2010; Hannaford et al. 2018). However, the precise spatio-temporal regulation of Actomyosin and the contribution to neuroblast asymmetric division are unclear.

In type I neuroblasts, at least five Actomyosin activity patterns occur at stereotypical times and locations along the cell cycle (**Figure 1A, MOV1**). The first of these patterns manifests as pulsatile non-muscle Myosin II (Myosin) activity during interphase/prophase driving cortical contractions (Oon and Prehoda 2021). Similar pulsatile contractions have been observed in many systems (Bement et al. 2024), and are driven by positive and negative feedback loops involving RhoGTPases, GAPs and GEFs in U2OS cells (Graessl et al. 2017), the *C. elegans* zygote (Michaux et al. 2018), starfish and frog embryos (Michaud et al. 2022) and *Drosophila* egg chambers (Jackson et al. 2024).

**Figure 1:**
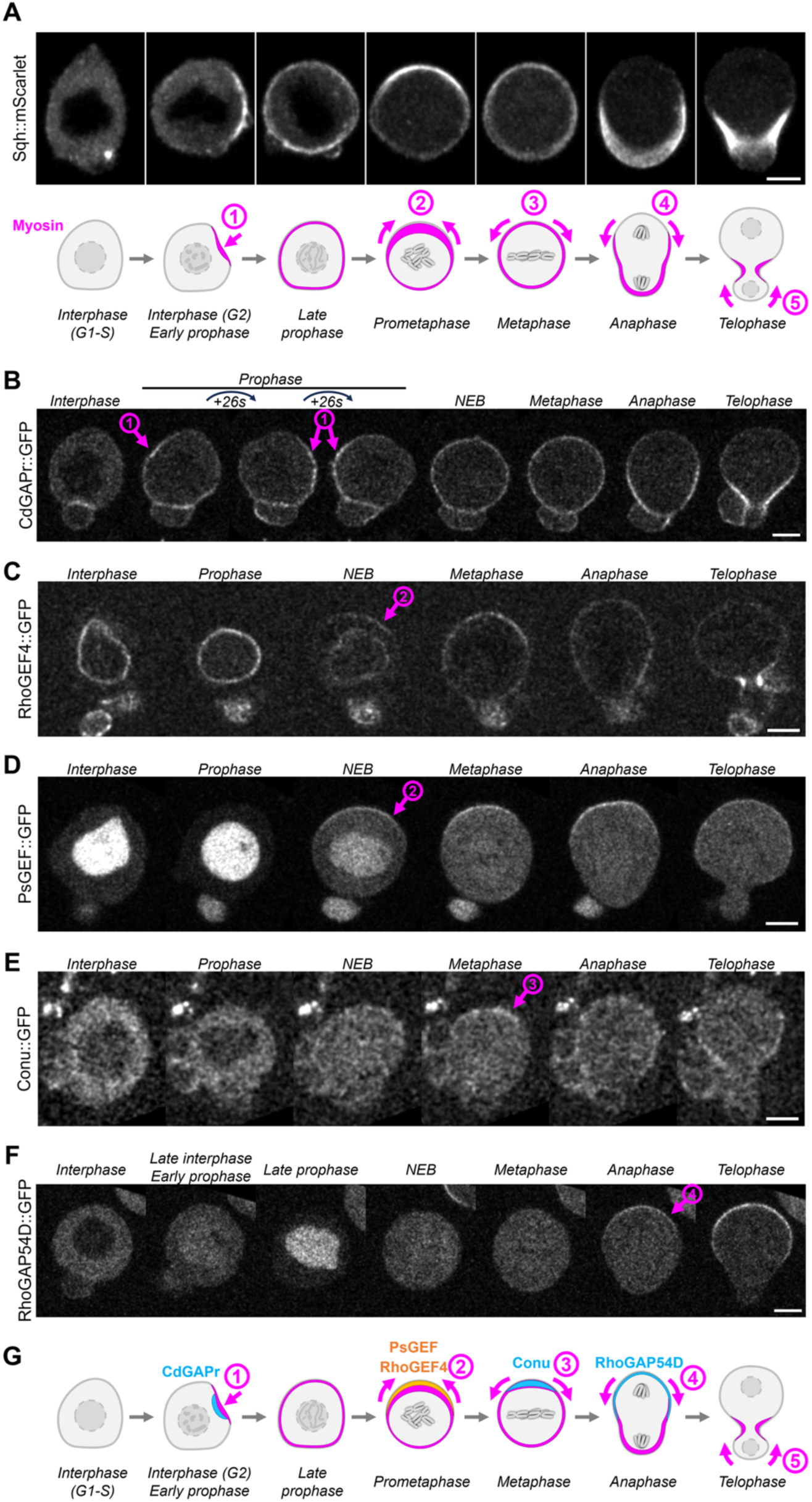
Cell cycle-dependent localisation of candidate RhoGAPs and RhoGEFs in type I neuroblasts. **A)** Reproduction of Myosin localisation during neuroblast asymmetric division: Myosin displays cell-cycle dependent localisation patterns. During interphase/prophase, localised cortical Myosin recruitment drives pulsatile contractions (1); at NEB, a basal-to-apical flow enriches Myosin at the apical pole (2); during metaphase, an apical-to-basal flow redistributes Myosin to the entire cortex (3); at anaphase, another apical-to-basal flow depletes Myosin form the apical pole (4); later in anaphase/telophase, a basal-to apical flow depletes Myosin from the basal pole (5). **B-G)** Localisation of candidates matching Myosin localisation patterns. *pulsatile contractions (1)*: CdGAPr is transiently enriched at the cortex during pulsatile contractions in interphase/prophase (arrow 1). **C-D)** *first basal to apical flow (2)*: RhoGEF4 **(C)** is recruited to the apical pole at NEB (arrow 2). PsGEF **(D)** is recruited to the apical pole at NEB (arrow 2). **E)** *first apical to basal flow* (3): Conu is recruited to the apical pole during metaphase (arrow3). **F)** *second apical-to-basal flow (4)*: RhoGAP54D is recruited to the apical pole during anaphase (arrow 4). Scale bar: 5 µm. **G)** Localisation of RhoGAPs and RhoGEFs suggesting their involvement in actomyosin flows during neuroblast divisions. Scale bars: 5 µm.

In neuroblasts, a second Myosin flow occurs at nuclear envelope breakdown (NEB). This flow is directed from the basal to the apical pole and coalesces polarity proteins into a defined apical cap (Oon and Prehoda 2019). This depends on CDK1 (Loyer et al. 2024), aPKC, that can affect the directionality of coalescence (Hannaford et al. 2018), and Pins (Tsankova et al. 2017). At NEB, Pins enriches Rho Kinase (Rok) apically (Tsankova et al. 2017), possibly driving this flow by locally enriching Myosin (Barros et al. 2003). This flow is reminiscent of the posterior-to-anterior Myosin flow polarising the *C. elegans* zygote, dependent on RHO-1 and its GEF ECT-2 (Pebble in flies) (Munro et al. 2004; Motegi and Sugimoto 2006).

A third flow, this time in apical-to-basal direction, redistributes apically enriched Myosin to homogenous cortical levels in metaphase (Tsankova et al. 2017). This is followed during anaphase by two final flows that are important for asymmetric cleavage furrow positioning and cell size asymmetry between the neuroblast and the GMC: Myosin is cleared from the apical pole earlier than the basal pole (Roubinet et al. 2017), making the apical pole transiently softer, thus allowing internal pressure to expand the apical membrane (Pham et al. 2019). Pins regulates both apical-to-basal flows: in metaphase, it recruits the kinase PKN to redistribute Rok and Myosin (Tsankova et al. 2017); during anaphase, it is necessary for early clearance of apical Myosin, but it is unclear how.

Given that in *pkn* mutant neuroblasts, cell size asymmetry is ultimately preserved (Tsankova et al. 2017), Pins must regulate cell size asymmetry through other yet to be identified mechanisms. Neuroblasts further monitor the orchestration of the anaphase flows positioning the cleavage furrow: in case the apical-to-basal flow at anaphase overshoots and shifts the cleavage furrow too far basally a correction mechanism involving a specific isoform of Pebble kicks in ensuring robust GMC size (Montembault et al. 2023).

Given the vast diversity of their protein domains (Tcherkezian and Lamarche-Vane 2007), RhoGAPs and RhoGEFs can orchestrate specific spatio-temporal regulation of Actomyosin downstream of various regulatory inputs. RhoGAPs and RHoGEFs are thus likely to participate in the orchestration of Actomyosin dynamics in neuroblasts. Given that some of the stereotypical Myosin patterns occur at precise cortical locations, we focussed on GAPs and GEFs with cortical localisation patterns matching Myosin dynamics. We systematically analysed the localisation of RhoGAPs and RhoGEFs in neuroblasts. We reveal that Par3/Baz and Pins recruit PsGEF, which in turn recruits RhoGAP54D to the apical pole specifically at anaphase. Impairing RhoGAP54D function perturbs apical clearing of Myosin in anaphase and results in neuroblasts and GMCs size defects. Thus, our study uncovers a mechanism downstream of apical polarity and Pins regulating neuroblast/GMC cell size asymmetry. We also identify an additional function of cytoplasmic RhoGAP54D in inhibiting pulsatile Myosin activity.

## Results

### Systematic analysis of RhoGAP and RhoGEF localisation during neuroblasts divisions

To identify regulators of the Myosin dynamics in neuroblasts (**Figure 1A, MOV1**), we used a collection of fly lines expressing GFP-tagged RhoGAPs and RhoGEFs at endogenous levels (di Pietro et al. 2023). We screened their localisation in type I larval neuroblasts, focussing on cortical recruitment to identifying candidates whose localisation suggests involvement in a specific Myosin behaviour. We analysed the 34 RhoGAPs and RhoGEFs predicted to be expressed in neuroblasts based on RNA-Seq data (Berger et al. 2012). We identified five candidates whose cortical localisation was compatible with a role in controlling specific Myosin behaviours in neuroblasts. All other observations are summarised in **Supplementary Table 1** and **Figure S1**.

The RhoGAP CdGAPr was the only candidate localising to the cortex during interphase and prophase. At this stage, Myosin drives pulsatile cortical contractions (**Figure 1A, arrow 1**). CdGAPr was transiently enriched at the cortex where these contractions took place (**Figure 1B, S2A,A’**, **MOV2**). CdGAPr remained cortical during metaphase and localised to the cleavage furrow at anaphase/telophase **(Figure 1B)**.

The RhoGEFs RhoGEF4 and PsGEF both showed nuclear localisation (the nuclear membrane and within the nucleus, respectively) in interphase and prophase. At NEB, both immediately appeared at the apical cortex (**Figure 1C-D, S2B-C’’’**), coinciding with the onset of the basal-to-apical flow at NEB (**Figure 1A, arrow 2**). Subsequently, cortical RhoGEF4 spread basally while remaining apically enriched, whereas PsGEF remained strictly apical. At anaphase/telophase, RhoGEF4 also localised to the cleavage furrow **(Figure 1C**, **MOV3)**. Intriguingly, the PsGEF signal transiently increased at the apical pole at anaphase (**Figure 1D, S2C’’**, **MOV4**), unlike other apically localised proteins such as Baz, whose signal quickly drops as it gets presumably diluted within the expanding apical cortex at anaphase (**Figure S2C’’**).

The RhoGAP Conundrum (Conu) was cytoplasmic from interphase to prometaphase and was recruited to the apical cortex during metaphase (**Figure 1E, S2D, MOV5**), coinciding with the onset of the apical-to-basal flow redistributing Myosin to the entire cortex at this stage (**Figure 1A, arrow3**). Conu was then recruited to the cleavage furrow at anaphase/telophase (**Figure 1E**).

Finally, RhoGAP54D was initially cytoplasmic in interphase but progressively recruited into the nucleus, until it was mostly nuclear by the end of prophase (**Figure 1F, S2E,E’)**. Nuclear recruitment of RhoGAP54D was slower than that of PsGEF (**Figure 1D, S2E’**) which was already nuclear a few minutes after nuclear reformation. Following NEB, RhoGAP54D stayed cytoplasmic but was recruited to the apical pole specifically at anaphase (**Figure 1F, S2,E’’, MOV6**). This localisation coincides with the apical-to-basal flow depleting Myosin from the apical pole at this stage (**Figure 1A, arrow3**).

Thus, our analysis of RhoGAPs and RhoGEFs uncovered proteins that could be regulators of some of the stereotype Myosin behaviours observed in neuroblasts (**Figure 1G**). We next sought to test their functional relevance using genetics and quantitative live cell imaging.

### RhoGAP54D drives apical Myosin depletion at anaphase promoting cell size asymmetry

We did not follow up on CdGAPr in this study but first tested whether RhoGEF4 and PsGEF control the basal-to-apical Myosin flow at NEB (**Figure 2A**). However, neither apical Myosin nor Baz enrichment, indicators of successful occurrence of this flow, were affected upon efficient *rhoGEF4* RNAi (**Figure S3A**) and in *psgef* null (Higuchi et al. 2009) mutants (**Figure 2B-C**). Of note, we observed that *psgef* mutant neuroblasts were significantly smaller than controls (14% smaller perimeter and 37% smaller volume, **Figure 2D**). We then tested whether Conu participates to the apical-to-basal flow at metaphase (**Figure 3A**). Again, efficient depletion of Conu (**Figure S3B**) did not significantly affect Myosin dynamics (**Figure 3B-C**). Thus, despite their localisation patterns, impairing the function of RhoGEF4, PsGEF and Conu does not significantly disrupt Myosin flows and Baz polarity at NEB and metaphase.

**Figure 2:**
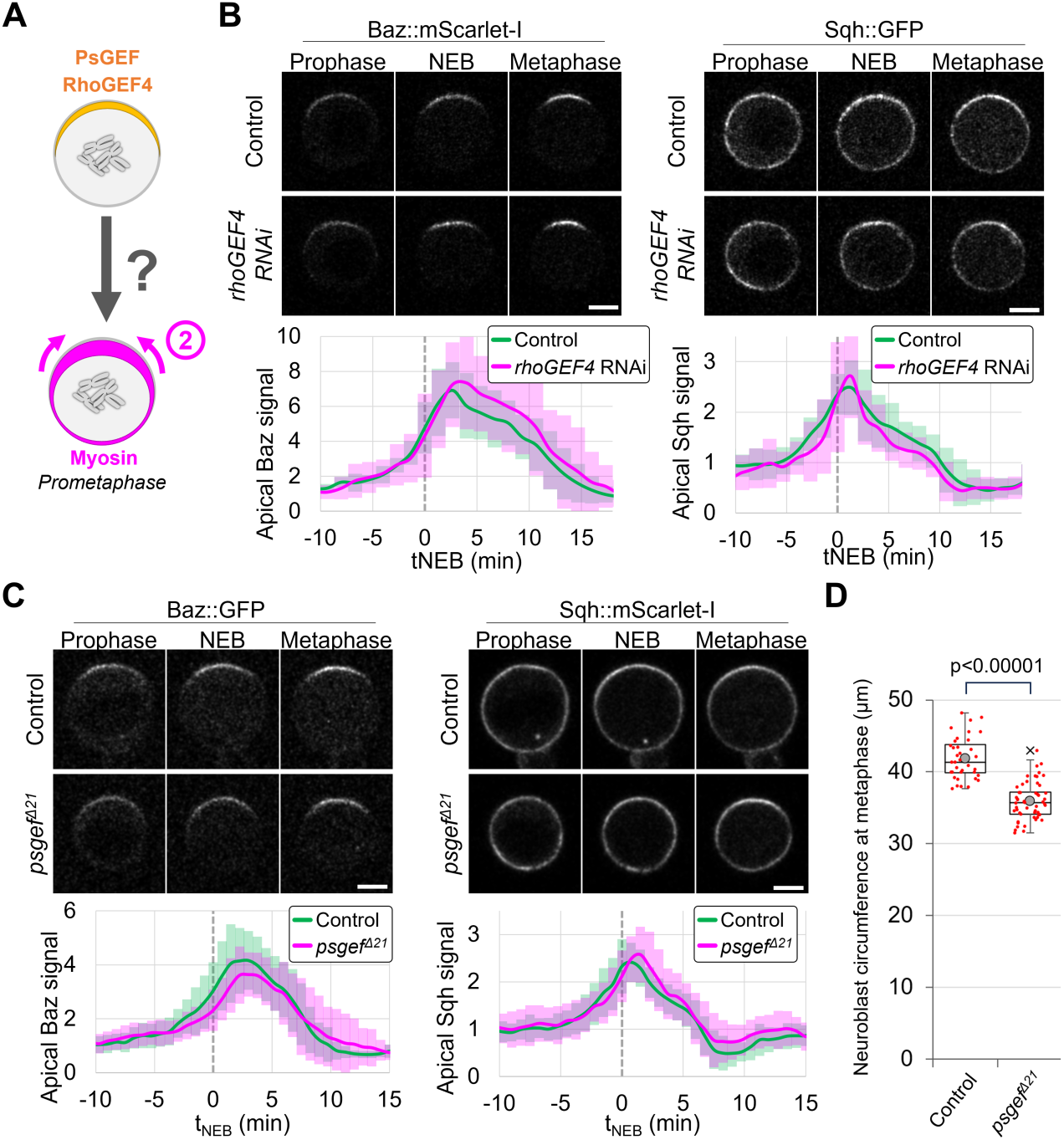
Interfering with RhoGEF4 or PsGEF function does not affect the basal-to apical flow at NEB. **A)** Hypothesis tested: The localisation of PsGEF and RhoGEF4 (orange) at the apical pole at NEB is involved in regulating the basal-to-apical Myosin (magenta) flow (arrows). **B)** Control and Neur-driven *rhoGEF4* RNAi neuroblasts (efficiency of rhoGEF4 RNAi has not been assessed). At NEB, Baz and Myosin increase in intensity at the apical pole in both controls (n=15) and RhoGEF4-depleted (n=11) neuroblasts. Bottom panels: intensity plot profiles of apical Baz and Sqh, normalised to the average signal across all control cells 10 to 15’ before NEB. Error bars: standard deviation. **C)** Control and *psgef^Δ21^* mutant neuroblasts. At NEB, Baz and Myosin increase in intensity at the apical pole in both controls (n=29) and *psgef^Δ21^* (n=20) neuroblasts. Bottom panels: intensity plot profiles of apical Baz and Sqh, normalised to the average signal across all control cells 10 to 15’ before NEB. 2 experiments. Error bars: standard deviation. **D)** The circumference of *psgef^Δ21^* neuroblasts at metaphase (35.9±2.6 µm, n=53) is smaller than in controls (41.9±2.9 µm, n=36). 3 experiments. Statistical test: two-tailed Mann– Whitney U test. Scale bars: 5 µm.

**Figure 3:**
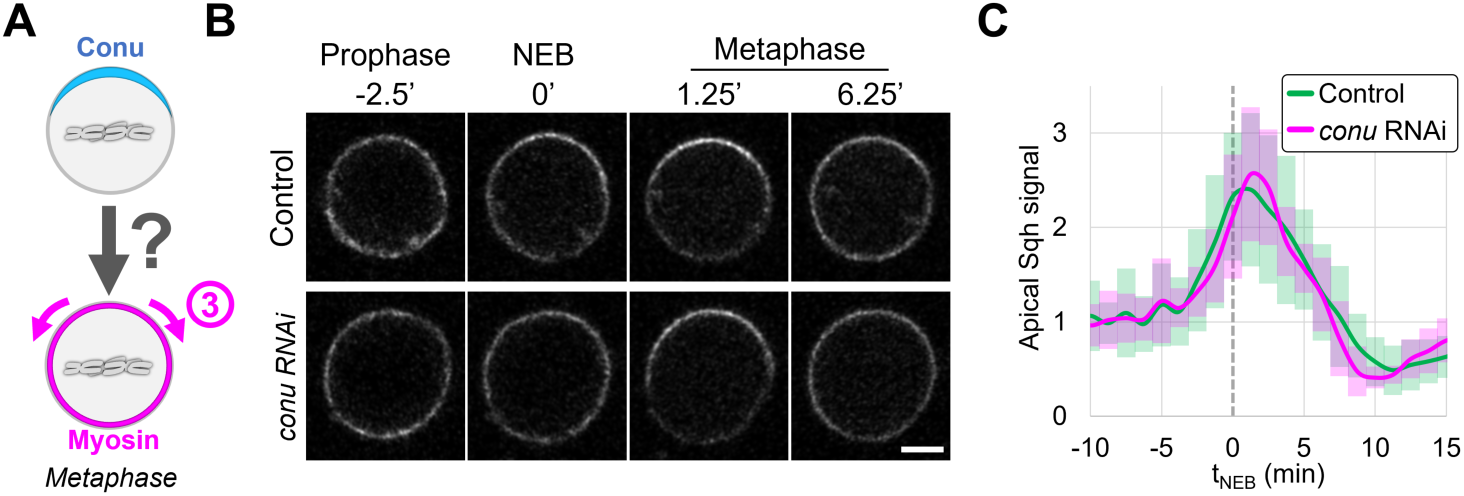
Loss of Conu does not affect Myosin redistribution at metaphase. **A)** Hypothesis tested: The localisation of Conu (blue) at the apical pole at metaphase regulates Myosin (magenta) redistribution (arrows). **B)** Control and Neur-driven *conu* RNAi neuroblasts. At metaphase, Myosin is initially enriched at the apical pole and then redistributed to the entire cortex in both controls and Conu-depleted neuroblasts. Scale bar: 5 µm. **C)** Intensity plot profile of apical Sqh in control (n=16) and Conu-depleted (n=21) neuroblasts, normalised to the average signal across all control cells 10 to 15’ before NEB. Error bars: standard deviation. 2 experiments.

We next turned our attention to RhoGAP54D, whose localisation suggests that it may be involved in the apical-to-basal flow at anaphase (**Figure 4A**). This flow depletes Myosin from the apical pole, which in turn promotes cell size asymmetry between the neuroblasts and its daughter cell (Roubinet et al. 2017). Remarkably, efficient depletion of RhoGAP54D by two different RNAi lines (**Figure S4**), individually or together, resulted in smaller neuroblasts (12% - 27% smaller perimeter and 31% - 61% smaller volume, **Figure 4B-C**). GMCs generated by RhoGAP54D-depleted neuroblasts were also abnormally large (18.4±2.5µm perimeter vs. 16.7±1.5µm in controls, **Figure 4D-E**). Thus, depleting RhoGAP54D affects neuroblast/GMC cell size asymmetry.

**Figure 4:**
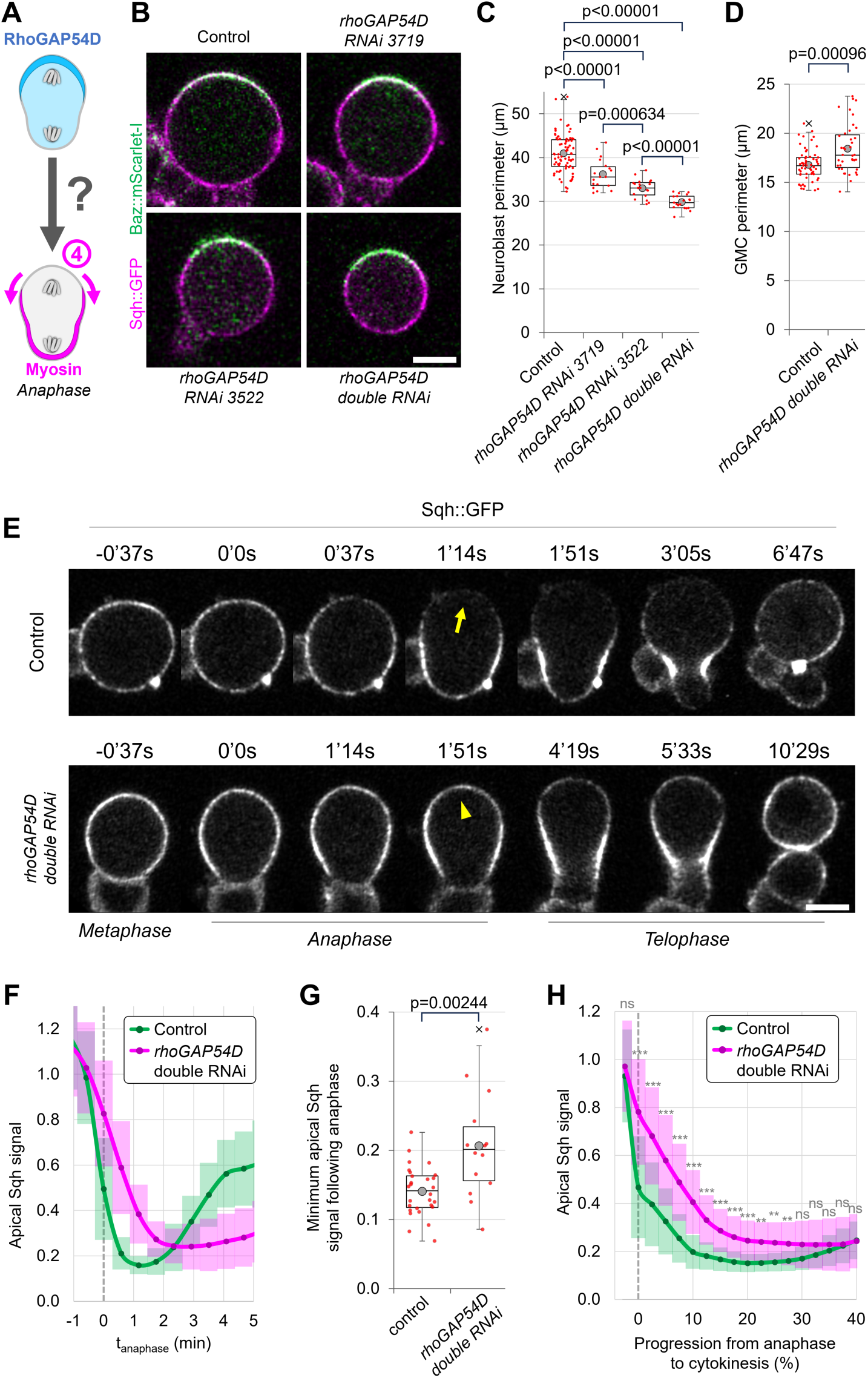
RhoGAP54D promotes apical Myosin depletion at anaphase and cell size asymmetry. **A)** Hypothesis tested: RhoGAP54D (blue) at the apical anaphase pole depletes Myosin(magenta) locally (arrows). **B)** Control and RhoGAP54D-depleted metaphase neuroblasts. RhoGAP54D depletion results in smaller cells. Scale bar: 5 µm. **C)** Neuroblast perimeter at metaphase. Controls: 40.9±4.3 µm, n=81; *rhoGAP54D RNAi 3719:* 36.2±3.3 µm, n=18; *rhoGAP54D RNAi 3522:* 33.0±2.1 µm, n=19; and *rhoGAP54D* double RNAi: 29.8±1.6 µm, n=19. 2 experiments. **D)** GMC perimeter in controls (16.8±1.6 µm, n=60) and *rhoGAP54D* double RNAi (18.4±2.1 µm, n=39). 2 experiments. **E)** Control and Neur-driven *rhoGAP54D* RNAi neuroblasts. Controls deplete Myosin efficiently apically (arrow) but not RhoGAP54D-depleted neuroblasts (arrowhead). **F)** Intensity plot profile of apical Sqh in control and RhoGAP54D-depleted neuroblasts, normalised to metaphase average intensity. **G)** Minimum apical Sqh levels during anaphase/telophase controls (n=28, 0.14±0.03) and RhoGAP54D-depleted neuroblasts (n=14, 0.21±0.7), normalised to metaphase average intensity. **H)** Intensity of apical Sqh in control and RhoGAP54D-depleted neuroblasts, normalised to metaphase average intensity, plotted along the percentage of progression from anaphase onset (0%) to cytokinesis completion (100%). Error bars: standard deviation. ***: p≤0.001; **: p≤0.01; ns: not significant. Statistical test: two-tailed Mann–Whitney U test. Scale bars: 5 µm.

We next analysed apical Myosin clearing at anaphase, a prerequisite for neuroblast/GMC cell size asymmetry. In controls, apical Myosin levels dropped to a minimum of 14±3% of their initial value at metaphase, whereas they only dropped to 21±7% upon RhoGAP54D-depletion (**Figure 4E-G, MOV7**). Apical clearing also appeared to be delayed (**Figure 4F**). To control that this was not due to the significantly longer anaphase of RhoGAP54D-depleted neuroblasts (**Figure 4E**, **S4B**), we plotted Myosin intensity along the percentage of progression from anaphase onset to cytokinesis. In controls Myosin levels had halved by anaphase onset (0%), whereas RhoGAP54D-depleted neuroblasts had to progress to 7.5% before dropping to this level (**Figure 4H**). Thus, the neuroblasts and GMCs size defects observed upon RhoGAP54D depletion are likely caused by late and incomplete apical Myosin clearing.

### RhoGAP54D inhibits pulsatile Myosin activity during early interphase and metaphase

While investigating the function of RhoGAP54D, we noticed that pulsatile Myosin-driven contractions in interphase/prophase (**Figure 1A, 5A**) occurred much earlier in RhoGAP54D-depleted neuroblasts than in controls (23.5±13.9’ and 16.8±9.0’ after cytokinesis in RNAi lines 3719 and 3522, respectively, vs 49±22’ in controls, **Figure 5A-B, MOV8**). Neuroblasts depleted of RhoGAP54D with both RNAi lines simultaneously stayed abnormally long in metaphase (**Figure S4C**), which allowed us to notice in two neuroblasts with particularly long metaphases (>30’) pulsatile cortical Myosin recruitment in metaphase. Contrary to interphase/prophase pulses, these pulses did not cause any visible contraction of the cortex (**Figure 5C**). To further investigate this, we arrested neuroblasts in metaphase. Controls displayed little variation of cortical Myosin (**Figure 5D-E**), whereas RhoGAP54D-depleted metaphase-arrested neuroblasts displayed stronger pulses of cortical Myosin that again did not appear to deform the cortex (**Figure 5D, MOV9**), resulting in a significant variation of cortical Myosin levels over time compared to controls (**Figure 5E-F**). We conclude that cytoplasmic RhoGAP54D inhibits pulsatile cortical activity of Myosin during early interphase and metaphase.

**Figure 5:**
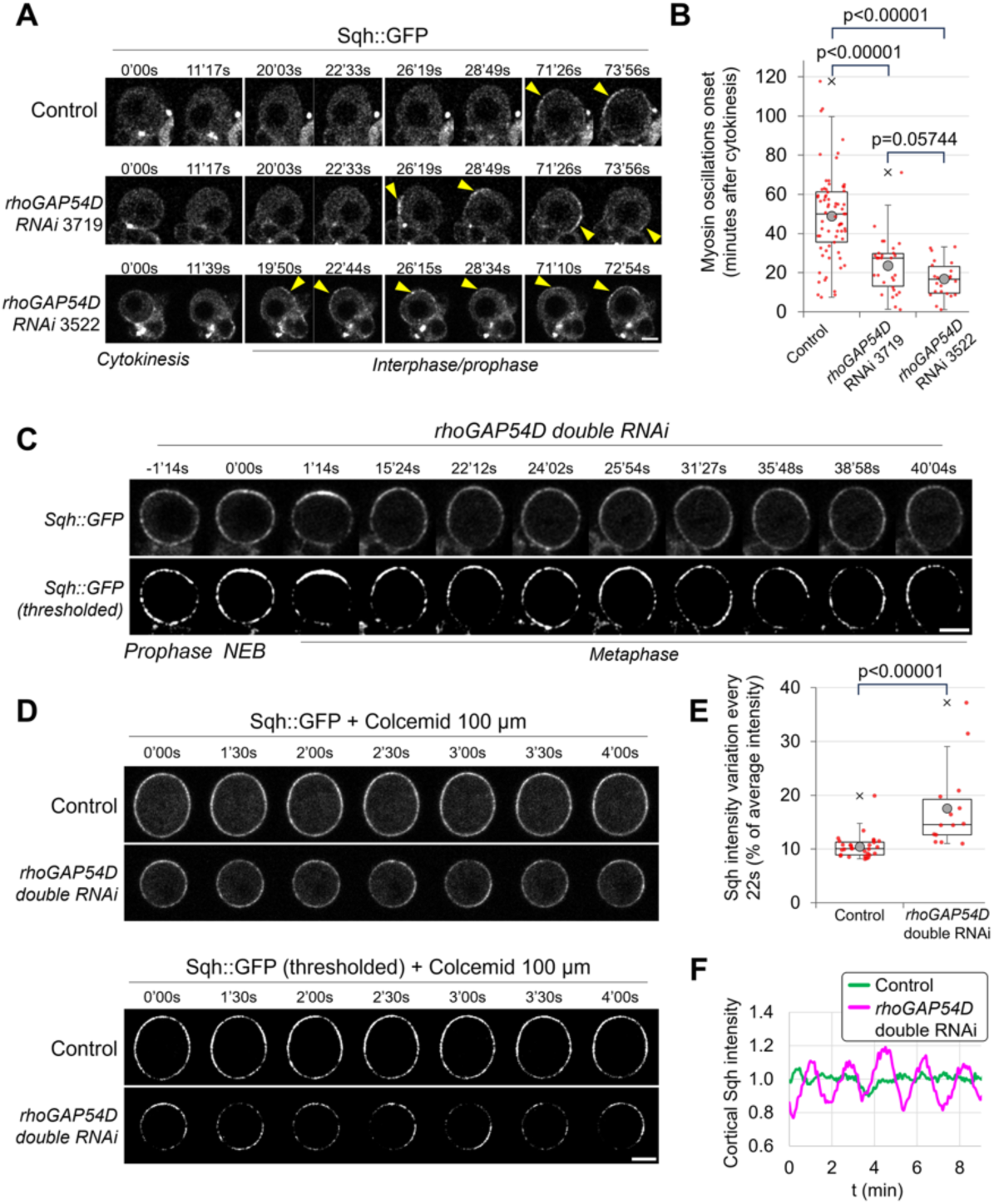
RhoGAP54D inhibits pulsatile Myosin contractions in early interphase and metaphase. **A)** Control and Neur-driven *rhoGAP54D* RNAi neuroblasts. Pulsatile cortical contractions (visualised by cortical recruitment of Myosin, yellow arrowheads) start earlier in RhoGAP54D-depleted neuroblasts. **B)** Quantification of the time of pulsatile contraction onset after cytokinesis: 49±22’ in controls (n=69), 24±14’ in *rhoGAP54D* RNAi 3719 (n=30), 17±9’ in *rhoGAP54D* RNAi 3522. 2 experiments. **C)** Cycling RhoGAP54D-depleted neuroblast with an abnormally long metaphase. Lower panels: the Sqh::GFP signal was thresholded to better visualise dynamic patches of local Myosin enrichment. **D)** Metaphase arrested control and RhoGAP54D-depleted neuroblasts. Lower panels: the Sqh::GFP signal was thresholded to better visualise dynamic patches of local Myosin enrichment. **E)** Average variation of cortical Myosin intensity (as a percentage of the average intensity over the imaging session) in metaphase-arrested neuroblasts. Myosin intensity varies by 10.4±2.2% every 22s in controls (n=29) and by 17.6±7.5% every 22s in RhoGAP54D-depleted neuroblasts (n=14). 2 experiments. **F)** Plot profiles of cortical Sqh::GFP intensity over time in the neuroblasts shown in panel **D** (normalised to average intensity over 8 minutes of imaging). Statistical tests: two-tailed Mann–Whitney U test. Scale bars: 5 µm.

### Mbo and PsGEF regulate the subcellular localisation of RhoGAP54D

We next investigated how the localisation of RhoGAP54D is regulated. A striking feature of RhoGAP54D localisation is its slow import into the nucleus (**Figure 1D, S2E’**). Interestingly, RhoGAP54D physically interacts with the nucleoporin Members Only (Mbo, (Giot et al. 2003), which regulates nuclear import of specific proteins (Uv et al. 2000). Mad, another nucleoporin-regulated protein (Colozza et al. 2011), also showed slow nuclear import (**Figure S2E’**). We therefore tested whether Mbo controls the nuclear import of RhoGAP54D. Likely due to pleiotropic effects, 27/30 Mbo depleted neuroblasts from larvae grown at 25°C did not divide within 1h30’. The 3 dividing neuroblasts, however, did not efficiently import RhoGAP54D into the nucleus (nuclear/cytoplasmic ratio of 1.2±0.2 at NEB, *vs* 9.7±3.7 in controls, **Figure 6A-B**). Mbo-depleted neuroblasts from larvae grown at 29°C (boosting UAS-dependent RNAi expression) did not divide at all and RhoGAP54D remained fully cytoplasmic in 13/21 neuroblasts (**Figure 6C**). Thus, Mbo regulates the nuclear import of RhoGAP54D.

**Figure 6:**
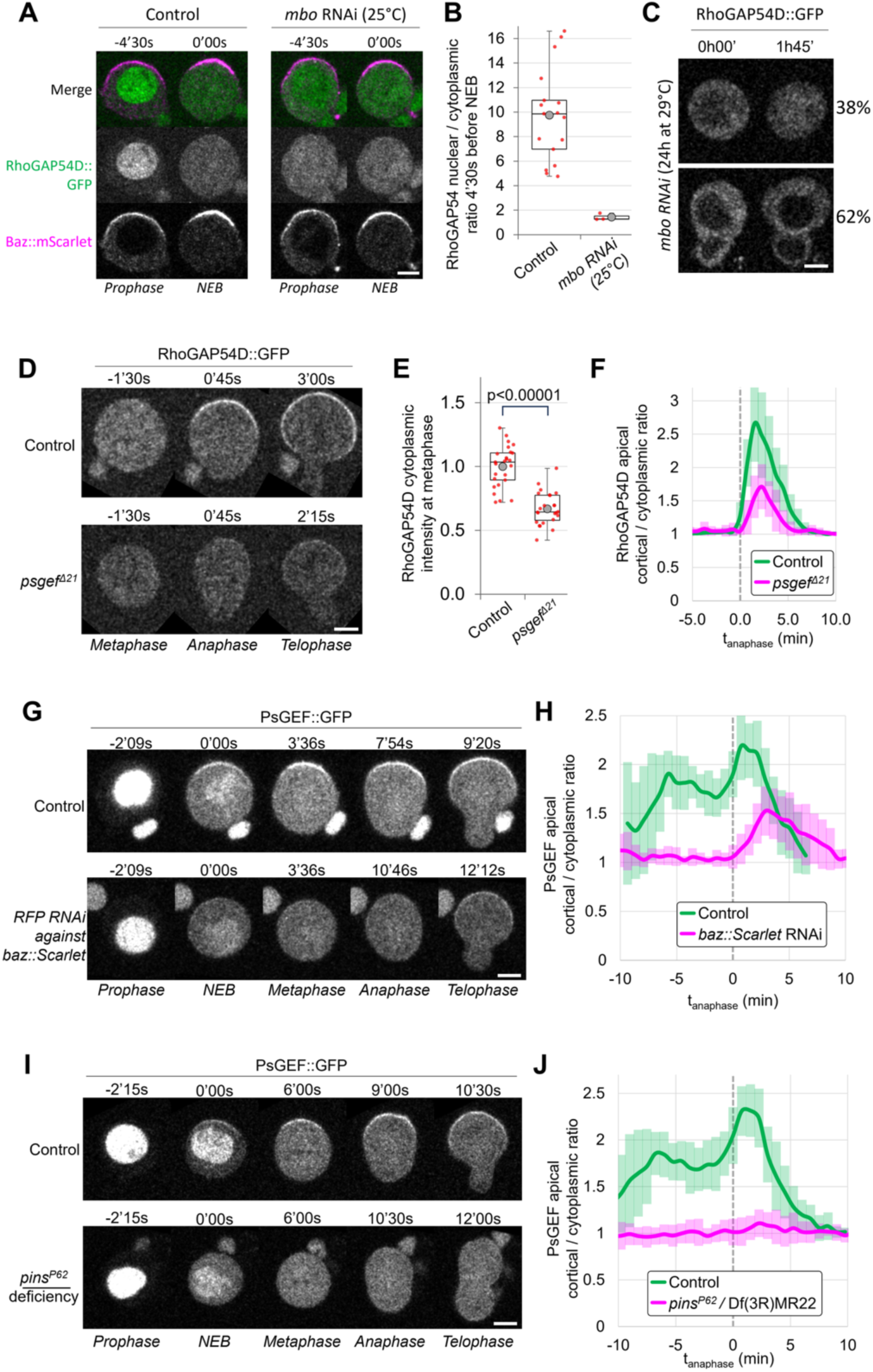
Mbo and PsGEF regulate the localisation of RhoGAP54D. **A)** Control and Neur-driven *mbo* RNAi neuroblasts (reared at 25°C). Just before NEB (visualised with Baz::mScarlet, magenta), RhoGAP54D::GFP (green) is mostly nuclear in controls whereas it is both in the nucleus and cytoplasm of *mbo* RNAi neuroblasts. **B)** Ratio of nuclear/cytoplasmic RhoGAP54D signal 4’30s before NEB in control (n=17) *vs mbo* RNAi (n=3) neuroblasts. 1 experiment. **C)** Neur-driven *mbo* RNAi neuroblasts from larvae grown at 29°C for 24 hours. *Mbo* RNAi Neuroblasts did not divide within 1h45’ of imaging (21/21). Some RhoGAP54D was imported into the nucleus during this time in only 38% (13/21) neuroblasts. 1 experiment. **D)** Control and *psgef^Δ21^* mutant neuroblasts. *psgef^Δ21^*: RhoGAP54D is 33±13% less abundant than in controls (determined by cytoplasmic levels at metaphase, **E**) and less efficiently recruited to the cortex (**F)** showing a maximal apical cortical/cytoplasmic ratio of 2.7±0.6 in controls (n=25), vs only 1.7±0.3 in *psgef^Δ21^* (n=24), 2 experiments. **G)** Control and Neur-driven *RFP* RNAi (depleting *baz::mScarlet*) neuroblasts. Upon Baz-depletion, PsGEF apical recruitment is abolished in metaphase and less efficient in anaphase/telophase. **H)** shows a maximal apical cortical/cytoplasmic ratio of 2.2±0.3 in controls (n=16), vs 1.5±0.3 upon Baz depletion (n=17) at anaphase/telophase, 2 experiments. **I,J)** PsGEF localisation in control and *pins* mutant neuroblasts. PsGEF apical recruitment is abolished in both metaphase and anaphase/telophase in *pins*. **J)** Plot profiles of apical cortical/cytoplasmic ratio for PsGEF in controls (n=17) vs *pins* (n=20) mutants, 2 experiments. Error bars: standard deviation. Statistical tests: two-tailed Mann–Whitney U test. Scale bars: 5 µm.

We next analysed the anaphase-specific apical recruitment of RhoGAP54D. Intriguingly, PsGEF, although being recruited to the apical pole at NEB, increases in intensity at anaphase (**Figure 1D, S2C’’-C’’’**). Furthermore, interfering with either PsGEF or RhoGAP54D leads to neuroblasts/GMCs size defects (**Figure 2C-D, 4B-C**). Thus, RhoGAP54D and PsGEF could function together in controlling neuroblast/GMC size asymmetry. The PsGEF isoforms expressed in neuroblasts (Berger et al. 2012) predominantly correspond to PsGEF-B, lacking the GEF domain (**Figure S5A,B**). Thus, rather than functioning as a GEF regulating Myosin activity, PsGEF could function to recruit RhoGAP54D to the cortex. For some reason, overall RhoGAP54D levels in the cytoplasm were 33% lower than in controls in *psgef* mutants (**Figure 6D-E**). Nonetheless, these lower RhoGAP54D levels were not efficiently recruited to the cortex (cortical/cytoplasmic ratio peaking at 1.7±0.3 in *psgef vs* 2.7±0.6 in controls, **Figure 6D,F, MOV10**). We conclude that anaphase specific recruitment of RhoGAP54D to the cortex depends on PsGEF.

### Pins recruits PsGEF to the apical neuroblast cortex in mitosis

Finally, we investigated factors that could regulate PsGEF localisation. The initial recruitment of PsGEF at NEB followed by a second round of recruitment at anaphase (**Figure 1D, S2C’’-C’’’**) could reflect involvement of two different regulators. We therefore tested the role of the two complexes defining apical polarity in neuroblasts: the Par and the Pins complex. The apical recruitment of PsGEF at metaphase was almost abolished upon *baz*-depletion (Par complex, achieved as in (Loyer et al. 2024) (**Figure S5C**) but was only delayed/weakened at anaphase (**Figure 6G-H, MOV11**). In contrast, the apical recruitment of PsGEF was abolished both at metaphase and at anaphase in *pins* mutants (**Figure 6I-J, MOV12**). In conclusion, the metaphase and anaphase apical recruitments of PsGEF are differentially regulated: at metaphase, both Baz and Pins are essential, whereas at anaphase, Baz is partially dispensable while Pins is essential.

## Discussion

To understand how Myosin regulation affects asymmetric neuroblast division, we aimed at identifying RhoGAPs and RhoGEFs expressed in *Drosophila* type 1 larval neuroblast that are involved in regulating the five stereotype patterns of Myosin activity described so far (**Figure 1A**). We identified five proteins whose cortical localisation matches four of the described Myosin patterns (**Figure 1**) and provide further insights into stereotypical Actomyosin regulation in neuroblasts (**Figure 7**).

**Figure 7:**
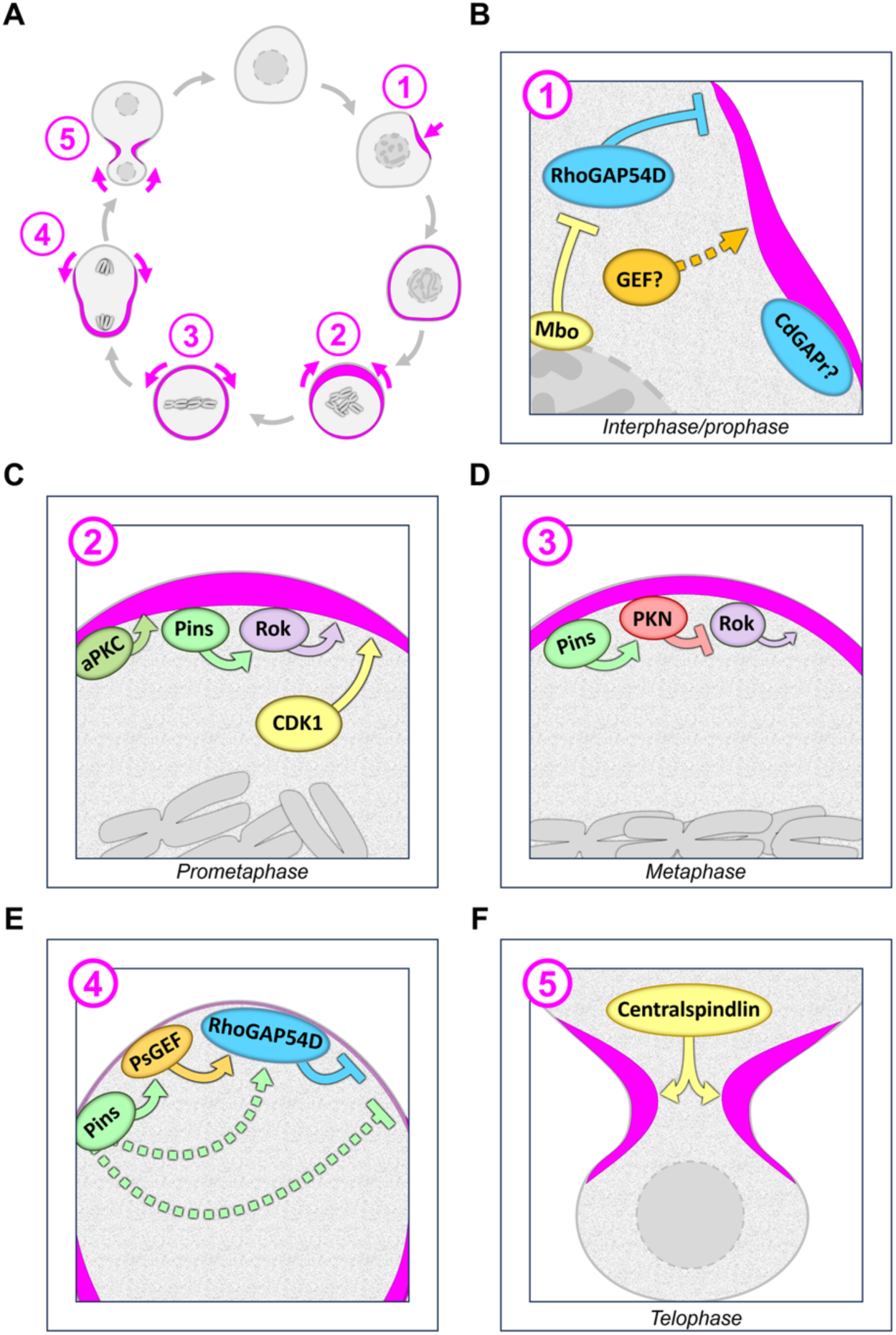
The regulation of Myosin dynamics along the neuroblasts cell cycle. See main text for details.

Pulsatile contractions constitute the first Myosin dynamics observed in late interphase/prophase and CdGAPr is the only candidate with a cortical localisation pattern corresponding to pulsatile contractions (**Figure 1E, 7B**). However, despite being cytoplasmic and not cortical at early interphase (**Figure 1F**), RhoGAP54D inhibits pulsatile contractions (**Figure 5**). RhoGAP54D’s function in regulating pulsatile contractions is likely controlled by nucleoporin-regulated nuclear import (**Figure 6A-C**): Pulsatile contractions do not occur at early interphase when RhoGAP54D is cytoplasmic but start later in interphase as RhoGAP54D is progressively sequestered in the nucleus. Finally, the pulses are largely inhibited by RhoGAP54D in metaphase (**Figure 5**) after full release from the nucleus (**Figure 1A, 1F**). This is reminiscent of the recent finding that nuclear sequestration and release of a RhoGAP/RhoGEF couple controls the timing of actomyosin pulses in *Drosophila* egg chambers (Jackson et al. 2024). We did not identify a cortical RhoGEF in interphase/prophase but, given that cytoplasmic RhoGAP54D controls pulses, cytoplasmic RhoGEFs may participate in this mechanism. Indeed, in adherent cells, mitotic cortical waves of active CDC42 encode positional information predicting the site of cleavage (Xiao et al. 2017). Whether the pulsatile contractions in neuroblasts are patterned and align with any aspect of asymmetric division has not been quantitatively studied yet.

Despite finding two RhoGEFs localising to the apical pole at NEB (**Figure 1C-D**), we did not identify any new regulator of the basal-to-apical flow occurring at NEB (**Figure 7C**). This flow is neither affected in *psgef* null mutants (**Figure 2A-B**), which is consistent with the fact that the PsGEF isoform expressed in neuroblasts likely does not have a GEF domain and thus may not regulate Myosin activity directly (**Figure S5A-B**), nor upon RNAi of *rhoGEF4*. Pins (Tsankova et al. 2017), aPKC (Hannaford et al. 2019) and CDK1 (Loyer et al. 2024) are currently known regulators of the basal-to-apical flow (**Figure 7C**), through mechanisms that remain to be elucidated. Similarly, Conu depletion does not affect Myosin redistribution during metaphase (**Fig 3, 7D**), which might not necessitate regulatory input by GAPs and GEFs as Pkn was proposed to directly phosphorylate Rok (**7D**). Interestingly, unlike Baz which appeared as smooth apical crescents in metaphase, Conu localised to apical membrane extensions originating from the apical pole (**Figure S1H**). Furthermore, apical membrane extensions are assembled by the Rac GTPase effectors SCAR and Arp2/3 (Cazzagon et al. 2023). Conundrum can somehow activate Rac (Neisch et al. 2013), and Mbc, a GEF for Rac (Geisbrecht et al. 2008), also localises to these extensions (**Figure S1I**). This suggests a function for Conu and Mbc in activating Rac at the apical pole in metaphase to organise apical membrane extensions.

A major finding of this study is that RhoGAP54D regulates neuroblast/GMC size asymmetry (**Figure 4**). RhoGAP54D localises to the apical pole exclusively at anaphase/telophase (**Figure 1F**), unlike any other reported protein localisation in neuroblasts. Our results show that Pins, PsGEF and RhoGAP54D function at anaphase to trigger the apical-to-basal flow at anaphase, a key driver of cell size asymmetry (Roubinet et al. 2017). Myosin flows in an apical-to-basal direction in a Pins-dependent manner both in metaphase (Tsankova et al. 2017) (**Figure 7D**) and anaphase (Roubinet et al. 2017) (**Figure 7E**) in what could appear as a single, continuous flow. However, the flows seem rather to be two distinct, differently regulated events as they can be temporally uncoupled: arresting neuroblasts in metaphase does not result in Myosin getting eventually cleared from the apical pole. Consistent with its apical localisation at metaphase, PKN redistributes Myosin to the entire cortex but is no longer apical at anaphase onset when Myosin is getting cleared (Tsankova et al. 2017). Pkn’s subcellular localisation is therefore less likely to trigger Myosin clearing and the flow at anaphase. In contrast, RhoGAP54D recruitment by Pins/PsGEF occurs at the right time and place to trigger Myosin depletion: apical GAP activity at anaphase provides a mechanistic explanation for locally downregulating Myosin to initiate the flow contributing to cell size asymmetry. This likely occurs *via* downregulation of Rho given that ARHGAP19, the human ortholog of RhoGAP54D, has a GAP activity toward RhoA, but not CDC42 or RAC (David et al. 2014). Importantly, unlike *pins* mutants, neither RhoGAP54D-depleted neuroblasts nor *psgef* mutants divide symmetrically. Therefore, Pins-dependent cell size asymmetry regulation likely involves additional mechanisms (**Figure 7E**). Finally, we did not find any apical RhoGEF or basal RhoGAP potentially regulating the final basal-to-apical flow (**Figure 7F**), consistent with the idea that this flow may be driven by a Myosin gradient at the lateral cortex (Roubinet et al. 2017).

The exact mechanism controlling anaphase-specific cortical recruitment of RhoGAP54D remains to be elucidated. Cortical recruitment of RhoGAP54D depends in part on PsGEF which is apically recruited by Baz and Pins, perhaps suggesting involvement of Inscuteable, which interacts with both Par and Pins polarity complexes (Schober et al. 1999; Wodarz et al. 1999; Schaefer et al. 2000). PsGEF is at the apical pole in metaphase (**Figure 1D**) but only promotes RhoGAP54D recruitment at anaphase. This is likely linked to the Pins-dependent second round of PsGEF apical recruitment at anaphase (**Figure 1D, 6I-J, S2C’’**): engaging in this mechanism may somehow enable PsGEF to recruit RhoGAP54D. PsGEF carries a C2 domain (**Figure S5B**) whose sequence most closely aligns with the C2 domain of the Synaptotagmin-like protein Bitesize, which binds phospholipids in a Ca^2+^-dependent manner (Pilot et al. 2006). Therefore, an increase of intracellular calcium at anaphase onset (Poenie et al. 1986) could drive the second round of PSGEF cortical recruitment. PsGEF also carries a domain of unknown function (**Figure S5B**). The uncharacterised protein CG15646 is almost entirely composed of this domain and interacts with Pins *in vitro* (Giot et al. 2003). Therefore, Pins may directly recruit PsGEF via this domain. In any case, another yet to be identified, likely Pins-dependent mechanism participates in RhoGAP54D recruitment, as RhoGAP54D localisation is not abolished in *psgef* mutants (**Figure 6 G,I**). CDK1 controls the cortical recruitment of RHoGAP54D’s human homolog ARHGAP19 (David et al. 2014; Marceaux et al. 2018) by phosphorylating a Serine conserved in RhoGAP54D (**Figure S6**), suggesting that the sudden drop of CDK1 activity at anaphase could trigger PsGEF-dependent and/or independent RhoGAP54D recruitment.

Human ARHGAP19 is involved in endometrial epithelium biology where its expression is regulated by microRNAs (Liang et al. 2021), at least in S2 cells RhoGAP54D expression is also reduced upon inhibition of miR-277 (Esslinger et al. 2013). Mutations in *ARHGAP19* have recently been identified in Charcot-Marie-Tooth patients and knockout models of ARHGAP19 in zebrafish and flies reveal that ARHGAP19 functions in promoting locomotor activity, pointing at loss of ARHGAP19 function as a cause of early-onset neuropathy (Dominik et al. 2024), yet the cell biological function of ARHGAP19 are not fully understood. Several characteristics of RhoGAP54D seem to be conserved in human ARHGAP19: like RhoGAP54D, ARHGAP19 physically interacts with nucleoporins (Müller et al. 2020), is nuclear in interphase and, most remarkably, stays in the cytoplasm in metaphase but gets recruited to the dividing cell poles of lymphocytes at anaphase (David et al. 2014). Another striking similarity between these orthologs is that they influence Actomyosin in metaphase despite being cytoplasmic: RhoGAP54D depletion causes Myosin pulses to continue in metaphase-arrested neuroblasts (**Figure 5D**) and ARHGAP19 depletion causes cell elongation in metaphase-arrested lymphocytes (David et al. 2014). Given these similarities, it would be interesting to test whether our findings about RhoGAP54D functions and regulation in the fly apply to ARHGAP19 to understand its biological functions and disease relevance.

## Acknowledgements

We thank Y. Bellaiche, E. Morais de Sa, M. Suzanne and HA. Müller for sharing flies. We thank the Dundee imaging facility for support. Stocks obtained from the Bloomington Drosophila Stock Center (NIH P40OD018537) were used in this study. This work was supported by grants from the BBSRC (BB/V001353/1 and BB/T017546/1). For the purpose of open access, the authors have applied a Creative Commons Attribution (CC BY) licence to any Author Accepted Manuscript version arising from this submission.

## Author contribution

JJ and NL conceived the study. NL designed all experiments and carried out most experiments. GL carried out experiments. NL and JJ analysed and interpreted the data and wrote the paper. JJ acquired funding and supervised the project.

## Declaration of interest

The authors declare no competing interest. GL’s current affiliation is Institute of Ophthalmology, University College London, London EC1V 9EL, United Kingdom.

Requests for further information and resources should be directed to and will be fulfilled by the lead contact, Jens Januschke.

## Supplementary figure legends

**Figure S1:**
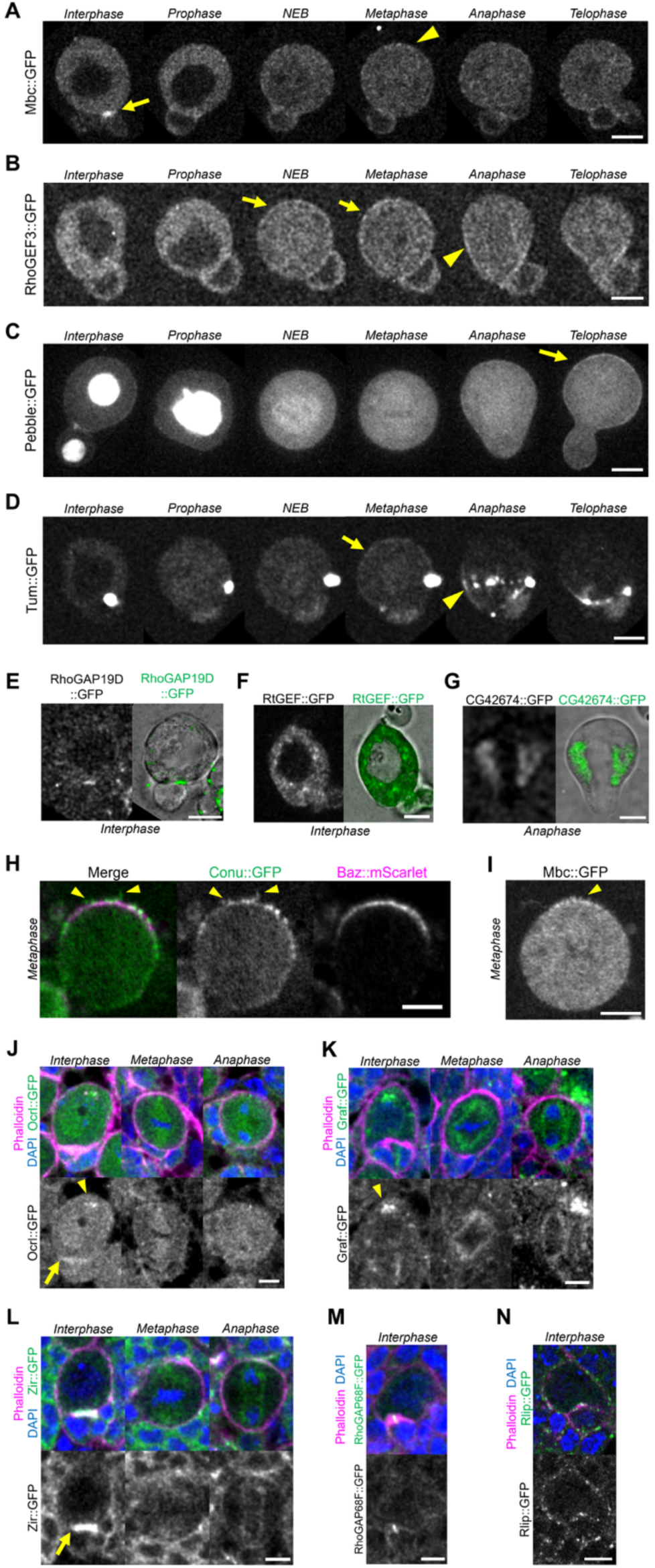
Localisation of further RhoGAPs and RhoGEFs in neuroblasts. **A-I)** Live neuroblasts in culture. **A)** Arrow: Mbc::GFP localisation at the midbody/neuroblast/GMC interface in interphase; arrowhead: cortical apical enrichment in metaphase. **B)** Arrows: cortical localisation of RhoGEF3::GFP. Arrowhead: cleavage furrow. **C)** Arrow: cortical Pebble::GFP at telophase. **D)** Arrow: cortical Tum at metaphase. Arrowhead: Tum localisation at the cytokinetic furrow and centralspindle. **E-F)** right panel: merge of the GFP signal (green) and transmitted light (gray). **E)** RhoGAP19D::GFP at the neuroblast/GMC interface. **F)**RtGEF::GFP on cytoplasmic punctae. **G)** CG42674::GFP on endomembrane surrounding the mitotic spindle at anaphase. **H-I)** Arrowheads: Conu::GFP and Mbc::GFP localisation at apical plasma membrane extensions in metaphase. **J-N)** Fixed neuroblasts in wholemount larval brains stained for DAPI (blue) and Phalloidin (Magenta). **J-L)** Arrows: Ocrl::GFP and ZirGFP at the neuroblast/GMC interface. Arrowheads: Ocrl::GFP and Gref::GFP at cytoplasmic punctae around the centrosome. **M**) RhoGAP68F::GFP at the midbody. **N)** Rlip at cortical punctae. Scale bars: 5µm

**Figure S2:**
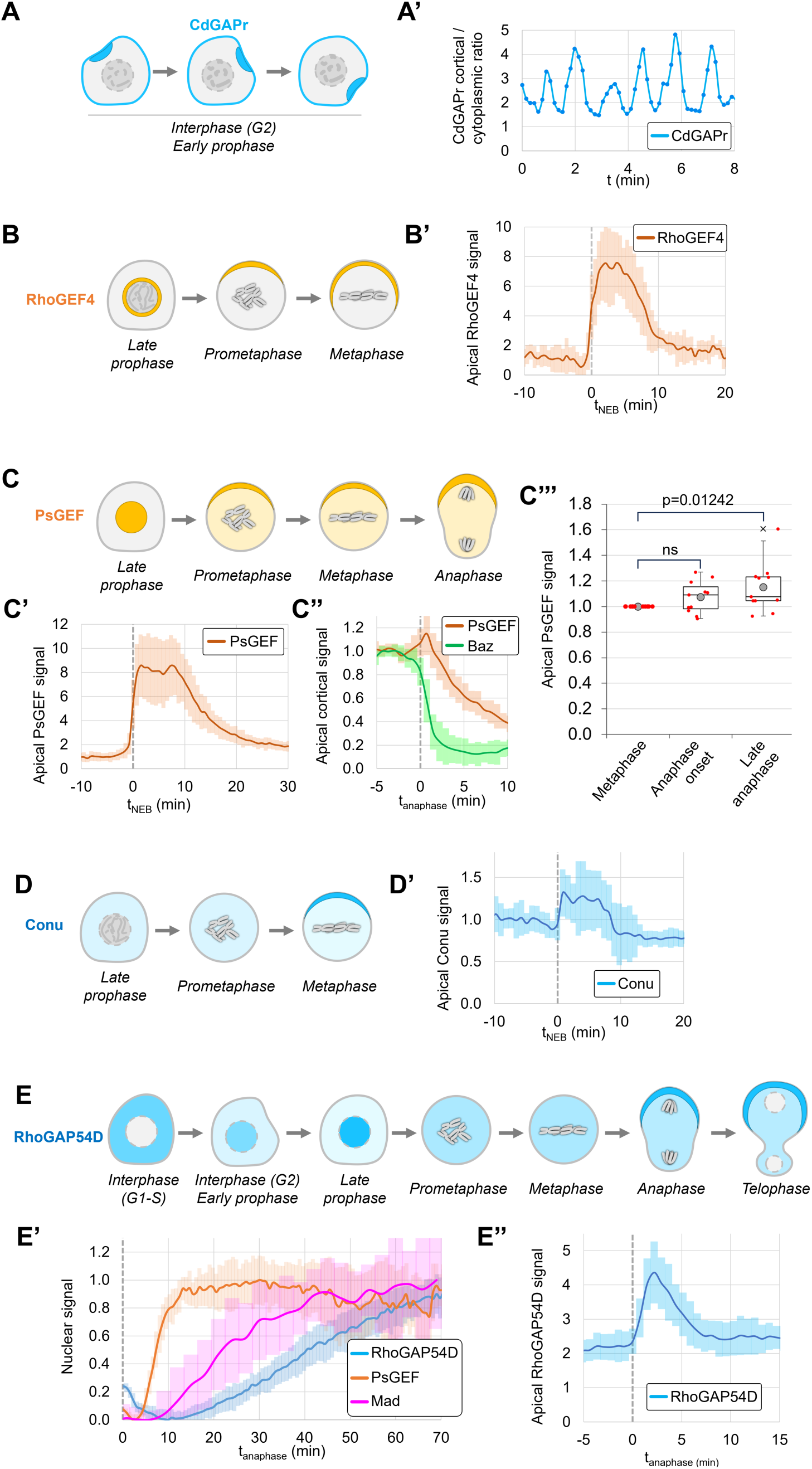
Time-resolved quantification of candidate RhoGAP and RhoGEF subcellular intensities. **A)** Schematic of Conu localisation. Conu is locally enriched at the cortex where pulsatile contractions occur. **A’)** Ratio of cortical / cytoplasmic Conu signal during interphase/prophase in one neuroblast. **B)** Schematic of RhoGEF4 localisation. RhoGEf4 relocalises from the nuclear membrane to the apical pole at NEB, then spreads further toward the basal pole during metaphase. **B’)**: apical RhoGEF4 signal during neuroblast division (n=9, 1 experiment), normalised to the average signal across all cells 5 to 10’ before NEB. t0: NEB. Error bars: standard deviation. **C)** PsGEF relocalises from the nucleus to the apical pole at NEB, then increases in intensity at the apical pole during anaphase. **C’)** Apical PsGEF signal normalised to the average signal across all cells 5 to 10’ before NEB. t0: NEB. **C’’)** Apical PsGEF and Baz signals normalised to the average signal across all cells 3 to 5’ before anaphase onset, t0: anaphase onset. Apical Baz levels drop very fast at anaphase onset, whereas PsGEF levels increase. **C’’’)** apical PsGEF signal at metaphase, anaphase onset (increase of 7±12%), and late anaphase (increase of 15±18%), individually normalised to the signal at metaphase. N=11, 1 experiment. Error bars: standard deviation. Statistical test: two-tailed Mann–Whitney U test. **D)** Conu relocalises from the cytoplasm to the apical pole during metaphase. **D’)** Apical Conu levels normalised to the average signal across all cells 5 to 10’ before NEB. t0: NEB. Error bars: standard deviation. N=9, 1 experiment. **E)** RhoGAP54D is cytoplasmic in early anaphase, slowly relocalises to the nucleus during interphase/prophase, is released into the cytoplasm at NEB and is recruited to the apical pole during anaphase/telophase. **E’)** Nuclear RhoGAP54D (n=30), PsGEF (n=11) and Mad (n=19) levels following anaphase, normalised to 0 for minimal levels and 1 for maximal levels. t0: anaphase. Error bars: standard deviation. **E’’)** Apical RhoGAP54D levels normalised to the average signal across all cells 5 to 10’ before NEB. N=15, 1 experiment.

**Figure S3.**
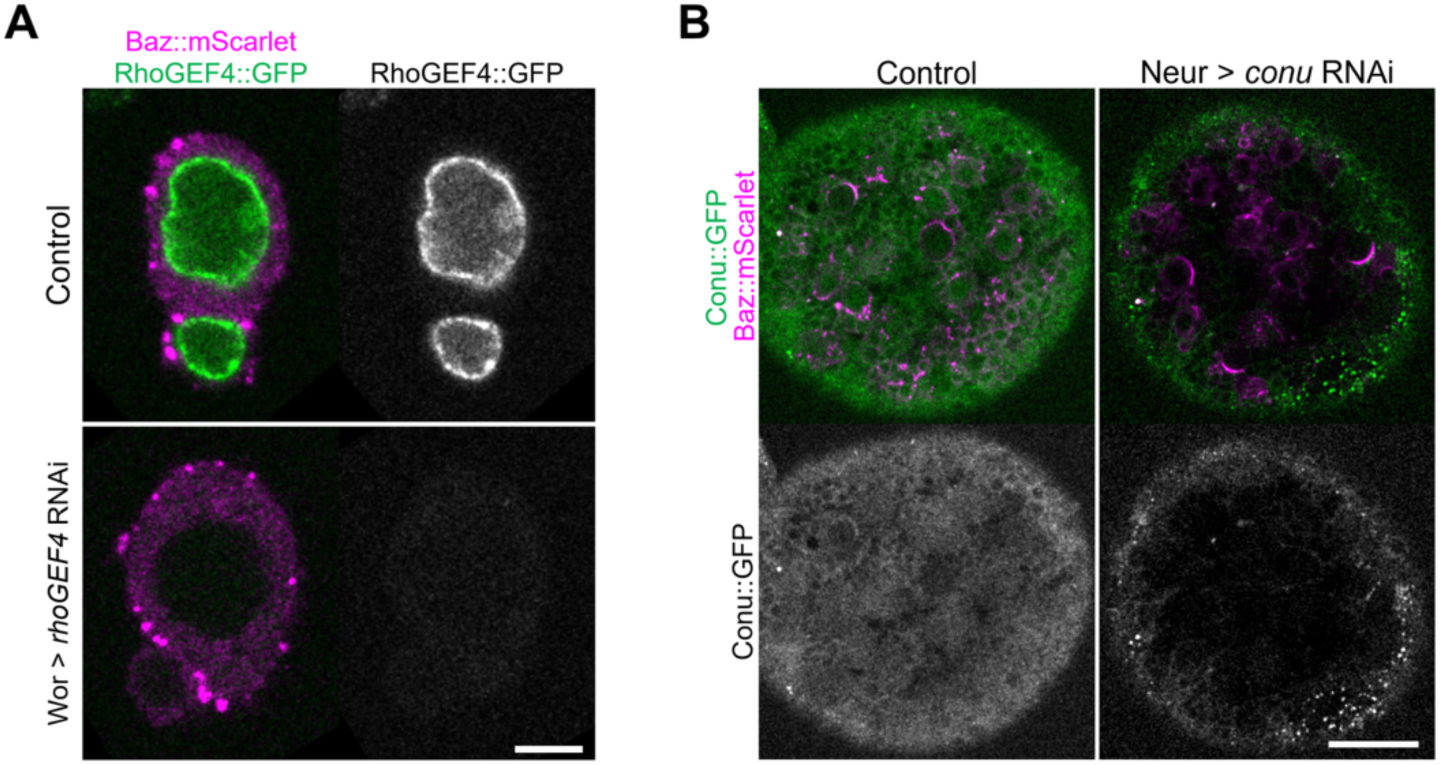
Efficient depletion of Conu and RhoGEF4 by RNAi. **A)** Isolated control and RhoGEF4-depleted neuroblasts. N=42 neuroblasts per condition, 1 experiment. Scale bar: 5 µm. **B)** Wholemount larval brains in control and Neur-driven *conu* RNAi. N=3 larval brains per condition, 1 experiment. Scale bar: 50 µm.

**Figure S4:**
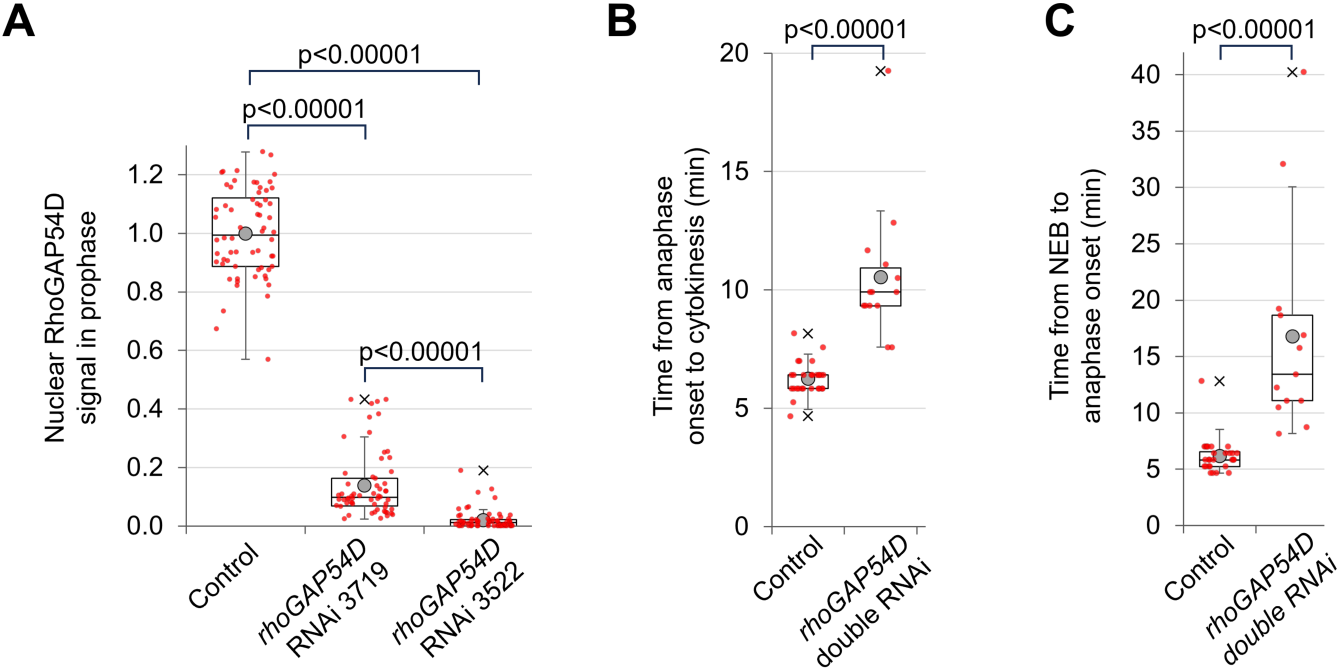
RhoGAP54D is efficiently depleted by RNAi which also affects the length of neuroblast mitosis. **A)** RhoGAP54D nuclear intensity 4’ before NEB in controls and Neur-driven *rhoGAP54D* RNAi, normalised to average value in controls. Controls: 1.00±0.15, n=64. RNAi line 3719: 0.14±0.11, n=60. RNAi line 3522: 0.02±0.03, n=67. 2 experiments. **B)** Time from anaphase onset to cytokinesis completion in controls (6.3±0.7’, n=24) and *rhoGAP54D* double RNAi (10.5±2.8’, n=14). 2 experiments. **C)** Time from NEB to anaphase onset in controls (6.2±1.5’, n=24) and *rhoGAP54D* double RNAi (17.8±9.1’, n=13). 2 experiments. Statistical test: two-tailed Mann– Whitney U test.

**Figure S5.**
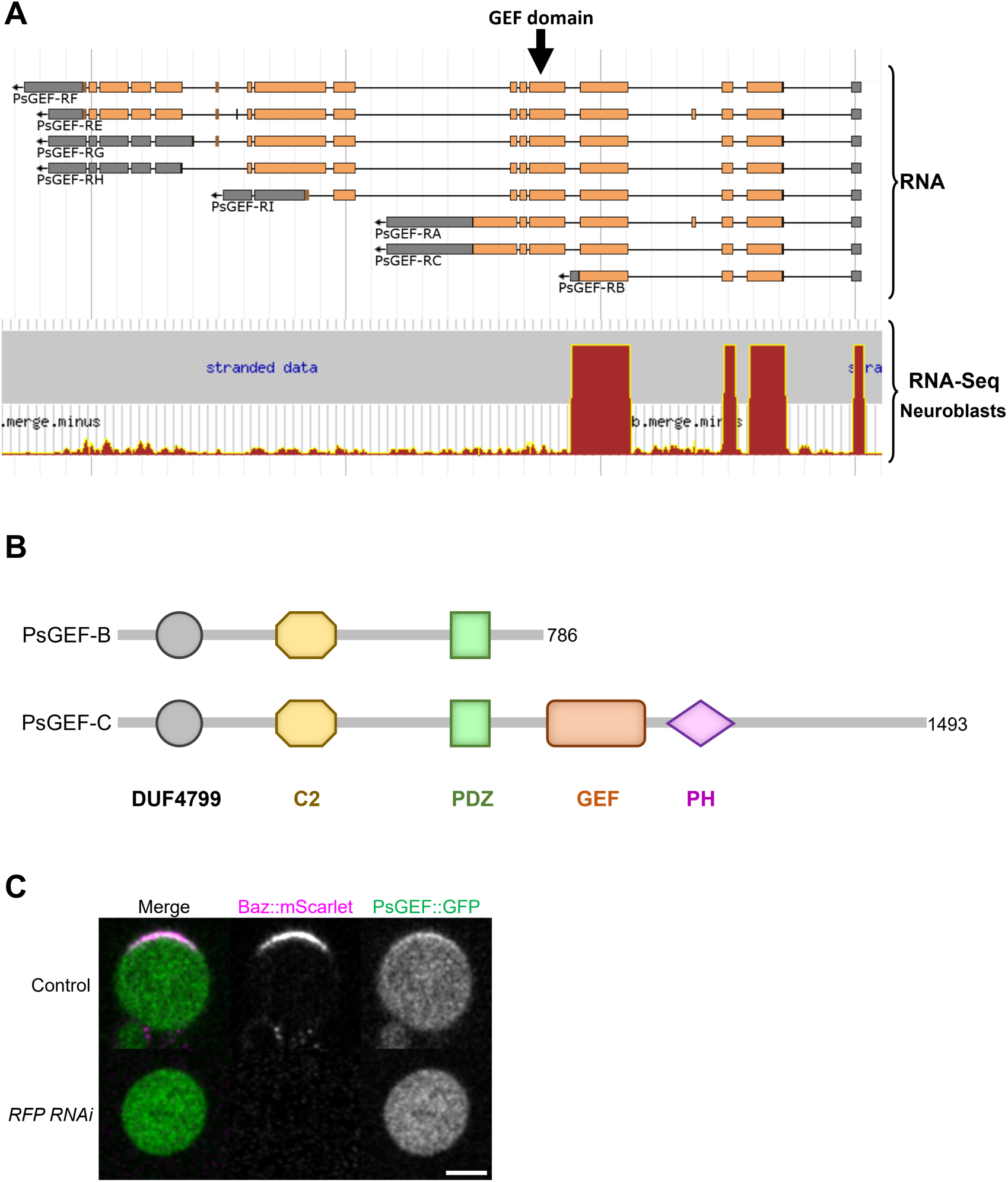
PsGEF-B lacks the GEF domain and is expressed in neuroblasts. **A)** Screenshot from Flybase, JBrowse. Top: “Reference Genome > RNA” track showing the spliced exons of alternative PsGEF isoforms. 5’ is right and 3’ is left as PsGEF is on the minus strand. Boxes: exons. Orange: coding region. Gray: untranslated region. The arrow points to the exon on which the GEF domain is encoded. Bottom: “Knoblich lab L3 CNS transcriptomes > L3 CNS neuroblast” track. Red bars: abundance of RNA sequences in neuroblasts. Display setting: “use linear signal scale”. Only the track corresponding to the minus strand is shown. **B)** Domains of the short isoform PsGEF-B and a longer isoform, PsGEF-C. **C)** Control and Neur-driven *RFP* RNAi neuroblasts in metaphase. RFP RNAi efficiently depletes mScarlet-tagged Baz (magenta). Scale bar: 5µm.

**Figure S6.**
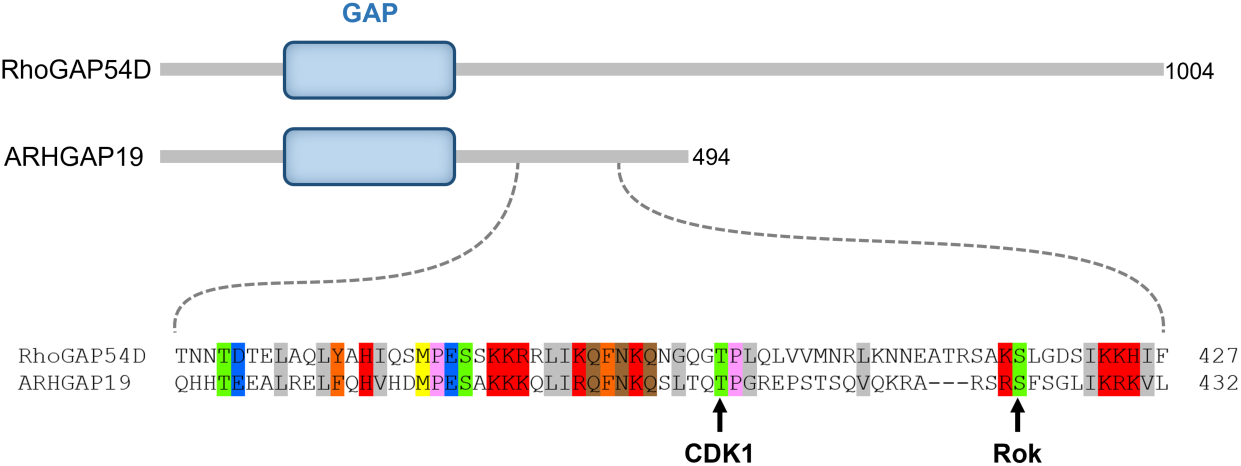
CDK1 and Rok phosphosites conservation between human ARHGAP19 and RhoGAP54D. Top: domain architecture of RhoGAP54D and ARHGAP19. Bottom: alignment of the RhoGAP54D and ARHGAP19 sequences around the ARHGAP19 residues phosphorylated by CDK1 and Rok (arrows).

## Movie legends

**MOV1: Stereotypical Myosin activity patterns along the neuroblast cell cycle.**

Neuroblast expressing Sqh::GFP. Arrows indicate stereotypic Myosin behaviour. 1) pulsatile Myosin contractions in late interphase/prophase (Oon and Prehoda 2021). 2) Basal to apical flow at NEB(Oon and Prehoda 2019). 3) Apical to basal flow at metaphase (Tsankova et al. 2017). 4) Apical to Basal flow at anaphase (Roubinet et al. 2017). 5) Basal to apical flow at anaphase (Roubinet et al. 2017). Neuroblast was imaged in primary cell culture, time stamp: mm:ss and scale bar 5 µm in this and all subsequent movies.

**MOV2: Representative movie of a neuroblast expressing CdGAPr::GFP.**

**MOV3: Representative movie of a neuroblast expressing RhoGEF4::GFP.**

**MOV4: Representative movie of a neuroblast expressing PsGEF::GFP.**

**MOV5: Representative movie of a neuroblast expressing Conu::GFP.**

**MOV6: Representative movie of a neuroblast expressing RhoGAP54D::GFP.**

**MOV7: RhoGAP54D depletion affects Myosin apical clearing at anaphase.**

Control and RhoGAP54D RNAi (double RNAi with lines 3719 and 3522) neuroblasts expressing Sqh::GFP side by side. Arrow: apical Myosin clearing defect. Arrowhead: an abnormally large GMC results from the division.

**MOV8: RhoGAP54D depletion causes pulsatile Myosin contractions to start earlier in the cell cycle.**

Control and RhoGAP54D RNAi (RNAi line 3719) neuroblasts expressing Sqh::GFP side by side. Neuroblasts synchronised at NEB. Arrowheads and arrows trace pulsatile Myosin contractions in the control and RhoGAP54D-depleted neuroblasts, respectively. t0: cytokinesis completion.

**MOV9: RhoGAP54D depletion causes pulsatile Myosin patterns in metaphase-arrested neuroblasts.**

Control and RhoGAP54D RNAi (double RNAi with lines 3719 and 3522) neuroblasts expressing Sqh::GFP side by side.

**MOV10: RhoGAP54D localisation in *psgef* mutants.**

Control and *psgef* mutant side by side. RhoGAP54D recruitment occurs with a delay and at lower levels (arrow) in the mutant. t0: anaphase onset.

**MOV11: PsGEF localisation upon Baz/Par-3 depletion.**

Control and *baz* depleted neuroblasts side by side. Cortical PsGEF recruitment is affected in metaphase (arrow), but less so in anaphase (arrowhead) upon *baz* depletion. t0: NEB.

**MOV12: PsGEF localisation in *pins* mutants.**

Control and *pins* mutant neuroblasts side by side. Cortical PsGEF recruitment is affected in metaphase (arrow) and in anaphase (arrowhead) in the mutant. t0: NEB.

## Material and methods

### Fly stocks and genetics

Flies were reared on standard corn meal food at 25 °C, except for RNAi-expressing larvae and their corresponding controls which were placed at 29 °C for 3 days before dissection. For the genotypes of the *Drosophila* lines used in each experiment, see **Supplementary Table 2**. For the origins of the stocks used, see **Supplementary Table 3**.

### Immunostainings

Larval brains were dissected in phosphate buffered saline (PBS) and fixed in 4% Formaldehyde (Sigma F8775) for 20 min at room temperature (RT), tissues were permeabilised in for 2 hr in PBS-Triton 0.1% (PBT) at RT. They were then incubated 1 hr at RT in a solution of DAPI 1/1000 (Sigma) and Phalloidin–Atto 647N 1/500 (Sigma) diluted in PBT, then for 1h in a solution of GFP-Booster Alexa Fluor 488 1/500 (ChromoTek), rinsed twice in PBT, washed for 10 minutes in PBT, rinsed twice in PBS, rinsed in 50% glycerol (Sigma 49781) and finally mounted between glass and coverslip in Vectashield (Vector Laboratories H-1000).

### Neuroblast primary culture

Every Figure except for the specified cases in Figure S1 shows live isolated neuroblasts. Every following reference to Schneider’s medium corresponds to Schneider’s medium (SLS-04– 351Q) supplemented with glucose (1 mg/ml), FCS 10% (ThermoFisher), fly extract 2.5% (DGRC) and insulin 0.07 mg/ml (ThermoFisher). Entire brains were dissected from L3 larvae in collagenase buffer (NaCl 8 mg/ml, KCl 0.2 mg/ml, NaH2PO4 0.05 mg/ml, NaHCO3 1mg/ml, D-Glucose 1mg/ml) and isolated from the surrounding imaginal discs. They were then incubated in 0.2 mg/mL collagenase (Sigma C0130-100MG) for 15 minutes, rinsed in collagenase buffer (800mg NaCl, 20mg KCl, 5mg NaH2PO4, 100mg NaHCO3, 100mg glucose in 100ml distilled water) and transferred into Schneider’s medium with 10mg/ml Fibrinogen (Januschke and Loyer 2020). A drop of 8 µl of Schneider’s medium with Fibrinogen containing the brains was pipetted onto a Poly-L-Lysin-coated glass dish. Brains were manually dissociated within the drop. The now isolated neuroblasts were left to settle for 5 minutes, after which clotting was induced by adding 1 µl of Thrombin 100 units/ml on 4 sides of the Schneider and Fibrinogen drop. Finally, clots were covered with 200 µl of Schneider’s medium. In experiments comparing control and RNAi or mutant neuroblasts, two clots with neuroblasts of different genotypes were prepared on the same dish.

### Live imaging and image processing

Isolated neuroblasts were imaged on a LEICA SP8 Stellaris confocal microscope equipped with an 86x water immersion objective (NA 1.20). A position was defined for each neuroblast in the Leica LAS X Navigator. Up to 120 neuroblasts, from one or two clots in the culture dish, were imaged every 20 seconds to 4 minutes on 1 to 7 confocal slices. A channel with the transmitted light was imaged in addition to any fluorophores expressed by neuroblasts. All images were processed and analysed using ImageJ (Schneider et al. 2012). A 2D gaussian blur with a sigma of 0.8 pixels was applied to all images displayed in Figures, but we performed intensity measurements on images without a gaussian blur. We corrected the drifting of movies with our custom AutoHyperstackReg macro (Januschke and Loyer 2020), based on the the TurboReg and MultiStackReg plugins (Thévenaz et al. 1998).

### Fluorescent signal measurement

Cytoplasmic or nuclear signal intensity was measured inside manually drawn polygonal shapes. Measurement of cortical signal intensity was performed using our custom rotating linescans macro (Januschke and Loyer 2020): briefly, the macro measures the signal along a band crossing the cortex in several orientation and retains the highest value found among all orientations. The background fluorescence was measured outside of neuroblasts and subtracted from any measured signal. Signal intensity was normalised as described in each Figure legend. Intensity profiles of individual neuroblasts were temporally aligned based on NEB or anaphase onset before being averaged. For the timing of NEB, we considered that NEB has started as soon as the Baz signal is no longer excluded from the nucleus. Anaphase onset was defined as the start of DNA segregation, visualised in the transmitted light channel.

### Cell size measurement

Neuroblasts size was measured by manually tracing the circumference of neuroblasts at metaphase. As neuroblasts are roughly spherical at this stage, the volume of neuroblasts was estimated with the formula V= (4/3) * π * (C/2π)^3^ where C is the circumference. For GMC size analysis, we used the transmitted light channel to manually draw the outlines of GMCs in interphase. As GMCs are not round at this stage, the volume cannot be calculated from these measurements.

### Quantitative data representation and statistical analysis

For every boxplot: cross: maximal and/or minimal outliers (beyond 1.5×interquartile range); grey circle: average; red dots: individual measurements; centre line, median; box limits, upper and lower quartiles; whiskers, 1.5×interquartile range. p Values were calculated using a non-parametric two-tailed Mann–Whitney U test in all cases.

References from tables (Buszczak et al. 2007; Proag et al. 2019; Hui et al. 2023; Li et al. 2024)

## References

Barros CS, Phelps CB, Brand AH. 2003. Drosophila nonmuscle myosin II promotes the asymmetric segregation of cell fate determinants by cortical exclusion rather than active transport. Dev Cell 5: 829–840.

Bement WM, Goryachev AB, Miller AL, von Dassow G. 2024. Patterning of the cell cortex by Rho GTPases. Nat Rev Mol Cell Biol 25: 290–308.

Berger C, Harzer H, Burkard TR, Steinmann J, van der Horst S, Laurenson AS, Novatchkova M, Reichert H, Knoblich JA. 2012. FACS purification and transcriptome analysis of drosophila neural stem cells reveals a role for Klumpfuss in self-renewal. Cell Rep 2: 407–418.

Bowman SK, Neumüller RA, Novatchkova M, Du Q, Knoblich JA. 2006. The Drosophila NuMA Homolog Mud regulates spindle orientation in asymmetric cell division. Developmental Cell 10: 731–742.

Broadus J, Doe CQ. 1997. Extrinsic cues, intrinsic cues and microfilaments regulate asymmetric protein localization in Drosophila neuroblasts. Curr Biol 7: 827–835.

Buszczak M, Paterno S, Lighthouse D, Bachman J, Planck J, Owen S, Skora AD, Nystul TG, Ohlstein B, Allen A et al. 2007. The carnegie protein trap library: a versatile tool for Drosophila developmental studies. Genetics 175: 1505–1531.

Cabernard C, Prehoda KE, Doe CQ. 2010. A spindle-independent cleavage furrow positioning pathway. Nature 467: 91–94.

Cazzagon G, Roubinet C, Baum B. 2023. Polarized SCAR and the Arp2/3 complex regulate apical cortical remodeling in asymmetrically dividing neuroblasts. iScience 26: 107129.

Cherfils J, Zeghouf M. 2013. Regulation of small GTPases by GEFs, GAPs, and GDIs. Physiol Rev 93: 269–309.

Colozza G, Montembault E, Quenerch’du E, Riparbelli MG, D’Avino PP, Callaini G. 2011. Drosophila nucleoporin Nup154 controls cell viability, proliferation and nuclear accumulation of Mad transcription factor. Tissue Cell 43: 254–261.

David MD, Petit D, Bertoglio J. 2014. The RhoGAP ARHGAP19 controls cytokinesis and chromosome segregation in T lymphocytes. J Cell Sci 127: 400–410.

Delgado MK, Cabernard C. 2020. Mechanical regulation of cell size, fate, and behavior during asymmetric cell division. Curr Opin Cell Biol 67: 9–16.

di Pietro F, Osswald M, De Las Heras JM, Cristo I, Lopez-Gay J, Wang Z, Pelletier S, Gaugue I, Leroy A, Martin C et al. 2023. Systematic analysis of RhoGEF/GAP localizations uncovers regulators of mechanosensing and junction formation during epithelial cell division. Curr Biol 33: 858–874 e857.

Dominik N, Efthymiou S, Record CJ, Miao X, Lin R, Parmar J, Scardamaglia A, Maroofian R, Aughey G, Wilson A et al. 2024. Biallelic variants in <em>ARHGAP19</em> cause a motor-predominant neuropathy with asymmetry and conduction slowing. medRxiv: 2024.2005.2010.24306768.

Esslinger SM, Schwalb B, Helfer S, Michalik KM, Witte H, Maier KC, Martin D, Michalke B, Tresch A, Cramer P et al. 2013. Drosophila miR-277 controls branched-chain amino acid catabolism and affects lifespan. RNA Biology 10: 1042–1056.

Gallaud E, Pham T, Cabernard C. 2017. Drosophila melanogaster Neuroblasts: A Model for Asymmetric Stem Cell Divisions. Results and problems in cell differentiation 61: 183–210.

Geisbrecht ER, Haralalka S, Swanson SK, Florens L, Washburn MP, Abmayr SM. 2008. Drosophila ELMO/CED-12 interacts with Myoblast city to direct myoblast fusion and ommatidial organization. Developmental Biology 314: 137–149.

Giot L, Bader JS, Brouwer C, Chaudhuri A, Kuang B, Li Y, Hao YL, Ooi CE, Godwin B, Vitols E et al. 2003. A protein interaction map of Drosophila melanogaster. Science 302: 1727–1736.

Graessl M, Koch J, Calderon A, Kamps D, Banerjee S, Mazel T, Schulze N, Jungkurth JK, Patwardhan R, Solouk D et al. 2017. An excitable Rho GTPase signaling network generates dynamic subcellular contraction patterns. J Cell Biol 216: 4271–4285.

Hannaford M, Loyer N, Tonelli F, Zoltner M, Januschke J. 2019. A chemical-genetics approach to study the role of atypical Protein Kinase C in Drosophila. Development 146: dev170589.

Hannaford MR, Ramat A, Loyer N, Januschke J. 2018. aPKC-mediated displacement and actomyosin-mediated retention polarize Miranda inDrosophilaneuroblasts. eLife 7: 166.

Higuchi N, Kohno K, Kadowaki T. 2009. Specific retention of the protostome-specific PsGEF may parallel with the evolution of mushroom bodies in insect and lophotrochozoan brains. BMC Biology 7: 21.

Hui J, Nakamura M, Dubrulle J, Parkhurst SM. 2023. Coordinated efforts of different actin filament populations are needed for optimal cell wound repair. Mol Biol Cell 34: ar15.

Jackson JA, Denk-Lobnig M, Kitzinger KA, Martin AC. 2024. Change in RhoGAP and RhoGEF availability drives transitions in cortical patterning and excitability in Drosophila. Curr Biol 34: 2132–2146 e2135.

Jaffe AB, Hall A. 2005. Rho GTPases: biochemistry and biology. Annu Rev Cell Dev Biol 21: 247–269.

Januschke J, Loyer N. 2020. Applications of Immobilization of Drosophila Tissues with Fibrin Clots for Live Imaging. J Vis Exp.

Li L, Zhang N, Beati SAH, De Las Heras Chanes J, di Pietro F, Bellaiche Y, Muller HJ, Grosshans J. 2024. Kinesin-1 patterns Par-1 and Rho signaling at the cortex of syncytial embryos of Drosophila. J Cell Biol 223.

Liang J, Li K, Chen K, Liang J, Qin T, He J, Shi S, Tan Q, Wang Z. 2021. Regulation of ARHGAP19 in the endometrial epithelium: a possible role in the establishment of uterine receptivity. Reprod Biol Endocrinol 19: 2.

Loyer N, Hogg EKJ, Shaw HG, Pasztor A, Murray DH, Findlay GM, Januschke J. 2024. A CDK1 phosphorylation site on Drosophila PAR-3 regulates neuroblast polarisation and sensory organ formation. Elife 13.

Loyer N, Januschke J. 2020. Where does asymmetry come from? Illustrating principles of polarity and asymmetry establishment in Drosophila neuroblasts. Curr Opin Cell Biol 62: 70–77.

Lu B, Ackerman L, Jan LY, Jan YN. 1999. Modes of protein movement that lead to the asymmetric localization of partner of Numb during Drosophila neuroblast division. Molecular cell 4: 883–891.

Marceaux C, Petit D, Bertoglio J, David MD. 2018. Phosphorylation of ARHGAP19 by CDK1 and ROCK regulates its subcellular localization and function during mitosis. J Cell Sci 131.

Michaud A, Leda M, Swider ZT, Kim S, He J, Landino J, Valley JR, Huisken J, Goryachev AB, von Dassow G et al. 2022. A versatile cortical pattern-forming circuit based on Rho, F-actin, Ect2, and RGA-3/4. J Cell Biol 221.

Michaux JB, Robin FB, McFadden WM, Munro EM. 2018. Excitable RhoA dynamics drive pulsed contractions in the early C. elegans embryo. J Cell Biol 217: 4230–4252.

Montembault E, Deduyer I, Claverie MC, Bouit L, Tourasse NJ, Dupuy D, McCusker D, Royou A. 2023. Two RhoGEF isoforms with distinct localisation control furrow position during asymmetric cell division. Nat Commun 14: 3209.

Motegi F, Sugimoto A. 2006. Sequential functioning of the ECT-2 RhoGEF, RHO-1 and CDC-42 establishes cell polarity in Caenorhabditis elegans embryos. Nat Cell Biol 8: 978–985.

Müller PM, Rademacher J, Bagshaw RD, Wortmann C, Barth C, van Unen J, Alp KM, Giudice G, Eccles RL, Heinrich LE et al. 2020. Systems analysis of RhoGEF and RhoGAP regulatory proteins reveals spatially organized RAC1 signalling from integrin adhesions. Nat Cell Biol 22: 498–511.

Munro E, Nance J, Priess JR. 2004. Cortical flows powered by asymmetrical contraction transport PAR proteins to establish and maintain anterior-posterior polarity in the early C. elegans embryo. Developmental Cell 7: 413–424.

Neisch AL, Formstecher E, Fehon RG. 2013. Conundrum, an ARHGAP18 orthologue, regulates RhoA and proliferation through interactions with Moesin. Mol Biol Cell 24: 1420–1433.

Oon CH, Prehoda KE. 2019. Asymmetric recruitment and actin-dependent cortical flows drive the neuroblast polarity cycle. eLife 8: 723.

Oon CH, Prehoda KE. 2021. Phases of cortical actomyosin dynamics coupled to the neuroblast polarity cycle. Elife 10.

Park HO, Bi E. 2007. Central roles of small GTPases in the development of cell polarity in yeast and beyond. Microbiol Mol Biol Rev 71: 48–96.

Parmentier ML, Woods D, Greig S, Phan PG, Radovic A, Bryant P, O’Kane CJ. 2000. Rapsynoid/partner of inscuteable controls asymmetric division of larval neuroblasts in Drosophila. J Neurosci 20: RC84.

Pham TT, Monnard A, Helenius J, Lund E, Lee N, Muller DJ, Cabernard C. 2019. Spatiotemporally Controlled Myosin Relocalization and Internal Pressure Generate Sibling Cell Size Asymmetry. iScience 13: 9–19.

Pilot F, Philippe JM, Lemmers C, Lecuit T. 2006. Spatial control of actin organization at adherens junctions by a synaptotagmin-like protein Btsz. Nature 442: 580–584.

Poenie M, Alderton J, Steinhardt R, Tsien R. 1986. Calcium Rises Abruptly and Briefly Throughout the Cell at the Onset of Anaphase. Science 233: 886–889.

Proag A, Monier B, Suzanne M. 2019. Physical and functional cell-matrix uncoupling in a developing tissue under tension. Development 146.

Riento K, Ridley AJ. 2003. Rocks: multifunctional kinases in cell behaviour. Nat Rev Mol Cell Biol 4: 446–456.

Roubinet C, Tsankova A, Pham TT, Monnard A, Caussinus E, Affolter M, Cabernard C. 2017. Spatio-temporally separated cortical flows and spindle geometry establish physical asymmetry in fly neural stem cells. Nature Communications 8: 237.

Schaefer M, Shevchenko A, Knoblich JA. 2000. A protein complex containing Inscuteable and the Galpha-binding protein Pins orients asymmetric cell divisions in Drosophila. Current biology: CB 10: 353–362.

Schneider CA, Rasband WS, Eliceiri KW. 2012. NIH Image to ImageJ: 25 years of image analysis. Nature methods 9: 671–675.

Schober M, Schaefer M, Knoblich JA. 1999. Bazooka recruits Inscuteable to orient asymmetric cell divisions in Drosophila neuroblasts. Nature 402: 548–551.

Staddon MF, Munro EM, Banerjee S. 2022. Pulsatile contractions and pattern formation in excitable actomyosin cortex. PLoS Comput Biol 18: e1009981.

Tcherkezian J, Lamarche-Vane N. 2007. Current knowledge of the large RhoGAP family of proteins. Biol Cell 99: 67–86.

Thévenaz P, Ruttimann UE, Unser M. 1998. A pyramid approach to subpixel registration based on intensity. IEEE Trans Image Process 7: 27–41.

Tsankova A, Pham TT, Garcia DS, Otte F, Cabernard C. 2017. Cell Polarity Regulates Biased Myosin Activity and Dynamics during Asymmetric Cell Division via Drosophila Rho Kinase and Protein Kinase N. Developmental Cell 42: 143–155.e145.

Uv AE, Roth P, Xylourgidis N, Wickberg A, Cantera R, Samakovlis C. 2000. members only encodes a Drosophila nucleoporin required for rel protein import and immune response activation. Genes Dev 14: 1945–1957.

Wodarz A, Ramrath A, Kuchinke U, Knust E. 1999. Bazooka provides an apical cue for Inscuteable localization in Drosophila neuroblasts. Nature 402: 544–547.

Xiao S, Tong C, Yang Y, Wu M. 2017. Mitotic Cortical Waves Predict Future Division Sites by Encoding Positional and Size Information. Dev Cell 43: 493–506 e493.

Yu F, Morin X, Cai Y, Yang X, Chia W. 2000. Analysis of partner of inscuteable, a novel player of Drosophila asymmetric divisions, reveals two distinct steps in inscuteable apical localization. Cell 100: 399–409.

